# Quantitative single cell mass spectrometry reveals the dynamics of plant natural product biosynthesis

**DOI:** 10.1101/2024.04.23.590720

**Authors:** Anh Hai Vu, Moonyoung Kang, Jens Wurlitzer, Sarah Heinicke, Chenxin Li, Joshua C. Wood, Veit Grabe, C. Robin Buell, Lorenzo Caputi, Sarah E. O’Connor

## Abstract

Plants produce an extraordinary array of complex natural products (specialized metabolites). Since the biosynthetic genes that are responsible for synthesis of these molecules are often localized to rare or distinct cell types, recently developed single cell RNA-sequencing (scRNA-seq) approaches have tremendous potential to resolve these complex pathways. In contrast, detection, identification, and quantification of metabolites in single cells has remained challenging. Here, we report a robust method for single cell mass spectrometry in which we rigorously characterize and quantify the concentrations of four classes of natural products in individual cells of leaf, root, and petal of the medicinal plant *Catharanthus roseus*. These single cell mass spectrometry datasets reveal information about the biosynthetic processes that cannot be determined from the corresponding scRNA-seq data alone, providing a highly resolved picture of natural product biosynthesis at cell-specific resolution.

## Introduction

Plants synthesize valuable natural products that are widely used in pharmaceutical, agrichemical, flavor, and fragrance industries. These complex molecules are synthesized by the sequential action of dedicated enzymes that exhibit highly regulated expression patterns among distinct cell types^1–3^. In one notable example, the ca. 40-enzyme biosynthetic pathway of the anti-cancer drug vinblastine (Fig. 1a), produced in the leaves of *Catharanthus roseus* (Madagascar periwinkle), occurs in three types of cells^4^: the first part of this alkaloid biosynthetic pathway occurs in internal phloem-associated parenchyma (IPAP) cells, the middle module of the pathway occurs in the epidermal cells, and finally, the last steps occur in the idioblast/laticifer cells.

**Figure 1.**
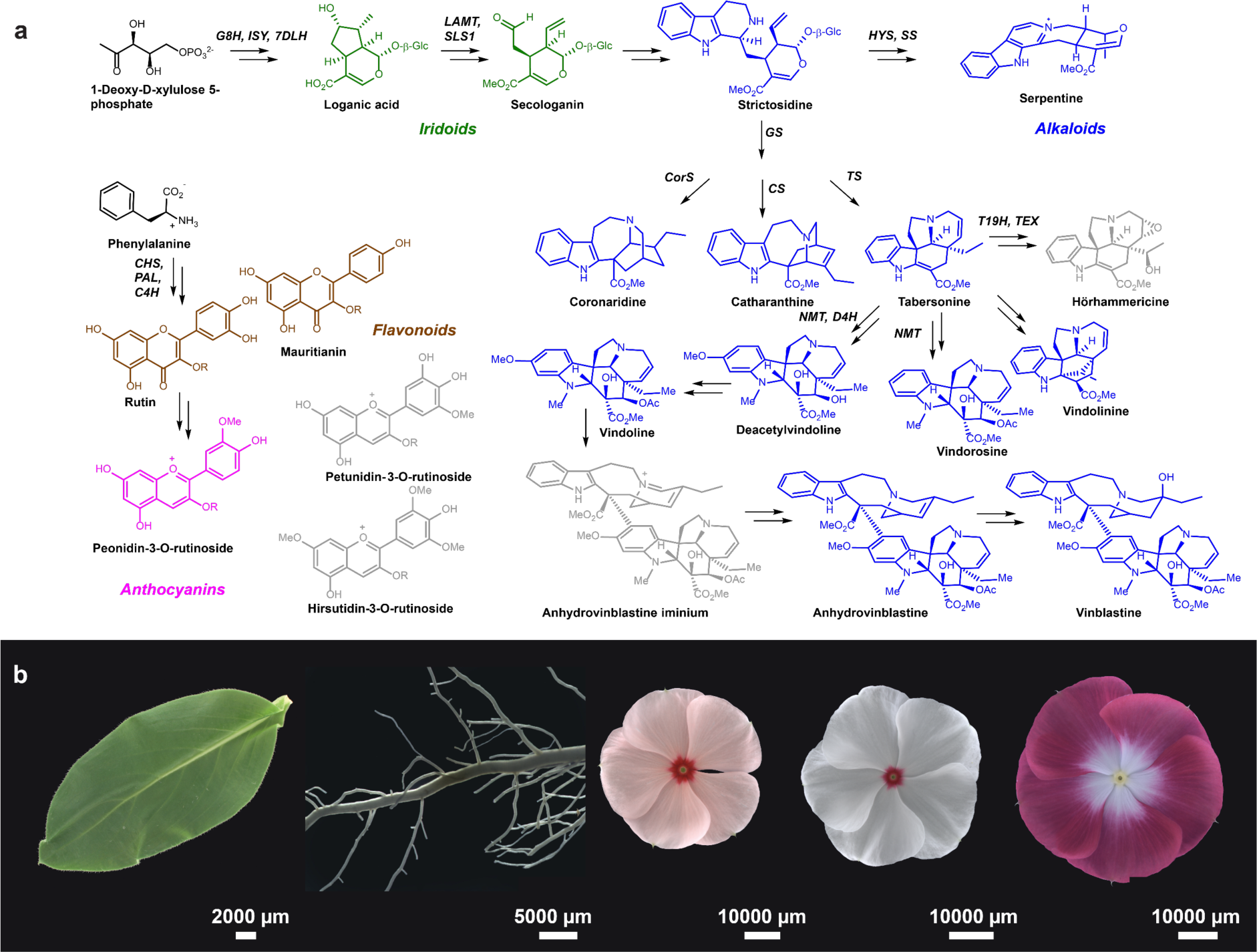
**a,** Simplified biosynthetic pathways of natural products produced in the medicinal plant *Catharanthus roseus*. Compounds in green are iridoid monoterpenes, which are also precursors for monoterpene indole alkaloids, shown in blue. Flavonoids are shown in brown and anthocyanins in pink. Compounds drawn in gray are mentioned in the main text but are not quantified with an authentic standard (See Supplementary Table 1 for definition of enzyme abbreviations). **b,** Photos of studied tissues (from left to right): Sunstorm Apricot (SA) leaf, SA root, SA flower, Little Bright Eyes (LBE) flower and Atlantis Burgundy Halo (ABH) flower.

The expression of different pathway genes in specific cell types makes plant specialized metabolism an ideal target for study using single cell RNA-sequencing (scRNA-seq) approaches. The cell-type specific expression profiles of the vinblastine biosynthetic genes of *C. roseus* leaves were recently revealed by scRNA-seq analyses^5, 6^. However, for scRNA-seq to be used effectively to understand natural product biosynthesis, the corresponding products and biosynthetic intermediates must also be mapped at the single cell level. Unfortunately, the applications for existing single cell mass spectrometry (scMS) methods in general (*e.g.* mass spectrometry imaging and live single cell mass spectrometry) are substantially limited by arduous sample preparation procedures and low throughput^7–9^. More importantly, these methods lack the possibility for chromatographic separation, which is required for both quantification and definitive structural identification^10^, essential information for understanding metabolic processes.

As part of a larger initiative to characterize the medicinally important alkaloids of *C. roseus* using scRNA-seq, we recently reported a preliminary scMS method for *C. roseus* leaf tissue that addressed these limitations^5^. With this approach, we could monitor a small number of alkaloids in single leaf cells using a cell-picking platform. Here, we show that this method can be broadly applied to cells derived from multiple tissues (leaf, root, and petal (Fig. 1b)), and can also be adapted to rigorously identify and quantify a range of metabolite classes (Fig. 1a) at an improved throughput of approximately 180 cells/day. Furthermore, using this method, we demonstrate how cell-specific localization patterns of alkaloid, phenylpropanoid, and monoterpene metabolite accumulation vary among organs. These scMS data also show that metabolites accumulate at highly variable levels within cell populations, with a minority of individual plant cells having alkaloids, iridoids and/or flavonoids at concentrations in excess of 100 millimolar. Overall, this scMS approach provides highly resolved profiles of how and where natural products are located at the cell-specific level, which, in combination with scRNA-seq, provides an improved foundation for gene discovery efforts, plant metabolic engineering and for understanding the function of natural products.

## Results

### Bulk tissue analysis of leaf, root and petal tissues

Before single cell analysis, bulk tissue analyses were performed using an Ultra High Performance Liquid Chromatography Q-Exactive Plus Orbitrap High Resolution Mass Spectrometry (UHPLC-HRMS) platform in the configuration used for single cell. The aim was to assess the chemical space of the plant tissues used for the single cell experiments and to validate instrument stability over three consecutive days of continuous measurements (Supplementary Table 2). Principal component analysis (PCA) of the processed data showed a clear separation of the tissues as expected (Supplementary Fig. 1). In these diluted bulk tissue samples, a total number of 1014 features with a molecular formula assigned with less than 2 ppm error was detected (Supplementary Data 1). This diluted sample, which yielded an MS signal comparable to what was observed in single cell analyses (see below), was used to validate the MS method.

The identities of bulk analysis features were first predicted based on their fragmentation spectra and library searches via SIRIUS^11–13^. The result from SIRIUS revealed the presence of 122 alkaloids, 66 flavonoids and anthocyanins, and 5 iridoids across all three tissues (Supplementary Data 2). We were able to confirm the identity of 18 metabolites using authentic standards and quantify 16 of these using external calibration (Supplementary Figs. 2-4, Supplementary Tables 3-4, Source File 1). These compounds included iridoids (loganic acid and secologanin), flavonoids (rutin and mauritianin), anthocyanin (peonidin-3-O-rutinoside) and a variety of monoterpene indole alkaloids (Fig. 1a, Supplementary Fig. 2, Supplementary Table 2). Mauritianin is a glycosylated form of kaempferol, whilst rutin is a quercetin glycoside and the occurrence of both the kaempferol and quercetin aglycones in *C. roseus* flowers is known^14^. The anthocyanin peonidin-3-*O*-rutinoside, for which a standard is also available, was detected in petals. Since the authentic standards for no other detectable anthocyanin in *C. roseus* were available, we made tentative assignments based on mass and fragmentation pattern of two additional anthocyanins, one (petunidin rutinoside-like) observed in all three cultivars, whilst the other (hirsutidin rutinoside-like) exclusively present in the ABH cultivar (Supplementary Fig. 2). Petunidin and hirsutidin scaffolds have been previously reported in *C. roseus* petals^15, 16^.

### The scMS workflow

A method for obtaining healthy and viable protoplasts for leaf, root and petal tissue of the Sunstorm Apricot (SA) cultivar was developed (see Methods). In addition, we examined petal tissue of Little Bright Eyes (LBE) and Atlantic Burgundy Halo (ABH) cultivars (Fig. 1b), since we anticipated that the petals of these differently colored cultivars would have different phenylpropanoid natural product profiles. Approximately 10,000 protoplasts were dispensed onto a microwell chip with cell-size micropores (50 µM) to capture single cells by gentle suction-induced sedimentation. Once situated in the micropores, cells were imaged by bright-field and fluorescence microscopy to record the size, morphology, and fluorescence (Supplementary Fig. 5b). In particular, we monitored the fluorescence signal as idioblast cells from *C. roseus* leaves display a characteristic blue fluorescence due to the accumulation of the alkaloid serpentine^17^, and we also monitored the presence of colored cells, which accumulated anthocyanins (Supplementary Fig. 5c). Cells were then collected into 96-well plates compatible with the autosampler of an UHPLC-HRMS system. Each 96-well contained 5 µL of 0.1% formic acid solution in water which resulted in lysis of each transferred protoplast by osmotic shock. Collection of each single cell takes approximately 13 seconds, providing an output of one 96-well plate in ca. 20 minutes. Plates were analyzed by UHPLC-HRMS after addition of internal standard and preparation of a pooled quality control sample. Chromatographic conditions were optimized for rapid analysis (7 minutes per run) allowing analysis of approximately 180 cells per day (Fig. 2). We previously showed that natural product profiles of *C. roseus* did not change substantially after protoplast isolation^5^.

**Figure 2.**
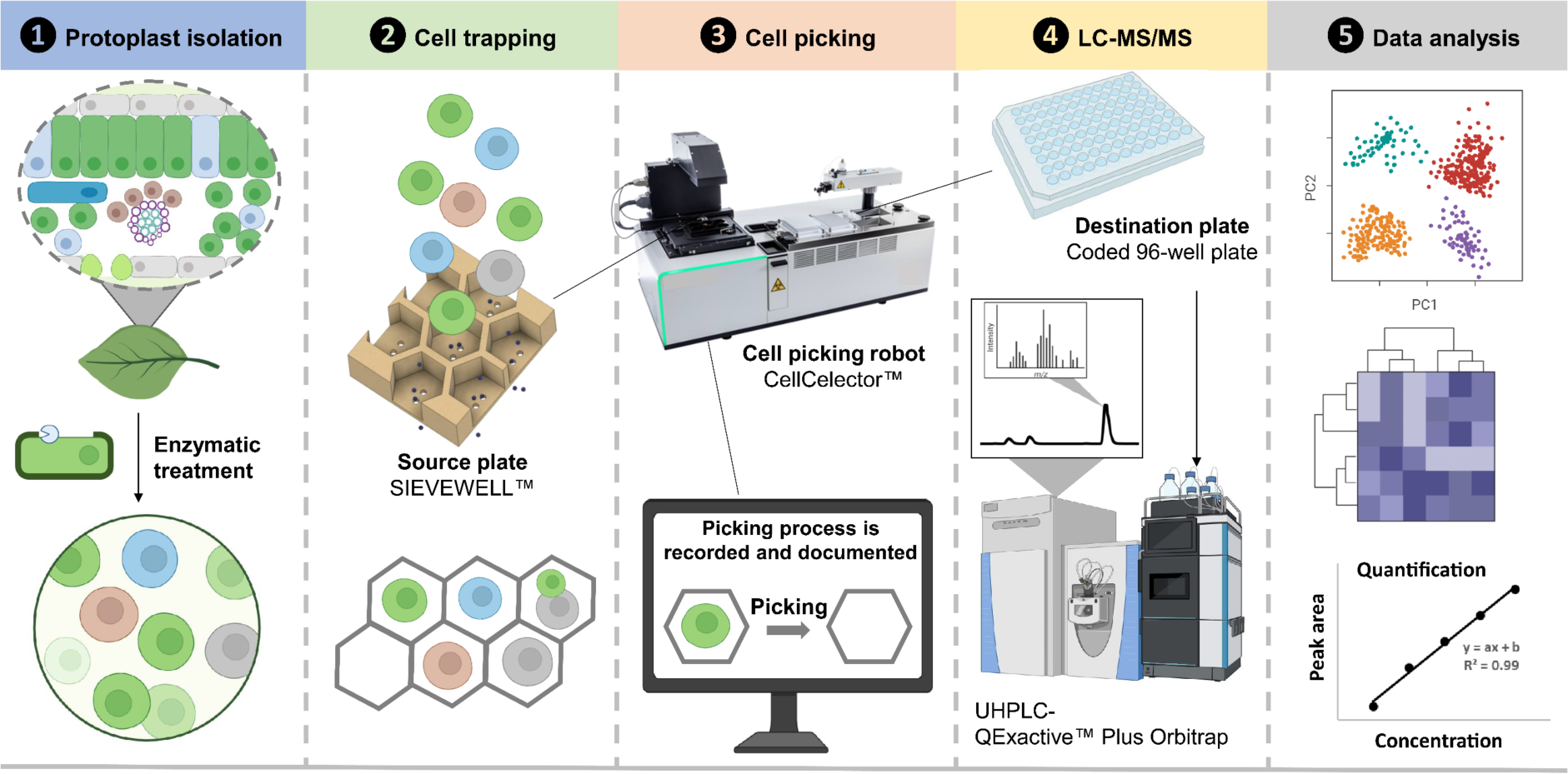
Workflow for single cell mass spectrometry (scMS) method described here. Protoplasts are isolated from leaf, root or petal, and trapped in the wells of a Sievewell plate. Individual cells are picked with a CellCelector™ robot and transferred to 96-well plates compatible with the autosampler of an LC-MS. Individual cells are then subjected to targeted and untargeted mass spectrometry.

### scMS of leaf-, root- and petal-derived single cells

Untargeted scMS was performed on approximately 200 cells from each of the five tissue samples (Supplementary Table 5, Supplementary Fig. 6-10). This untargeted metabolomic data processing pipeline extracted approximately 1000 features having a chemical formula assignment with less than 2 ppm error (Supplementary Table 5, Supplementary Data 3). The chromatography method was optimized for detection of alkaloid, iridoid, and phenylpropanoid natural products; primary metabolites such as amino acids and lipids were not captured in this analysis. The number of robust features identified in each analyzed cell was estimated from 10 randomly selected cells from each dataset; in these representative cells, the number of features varied between 10 and 280, reflecting the difference in the natural product content among the cell population (Supplementary Table 5). Hierarchical clustering analysis of the untargeted dataset from leaf using a subset of 39 features that could be confidently assigned as iridoid, phenylpropanoid, or alkaloid was performed (Fig. 3). Immediately apparent was the small number of cells that were highly enriched in alkaloids, whereas a larger number of cells accumulated flavonoids, secologanin, and a lower concentration of alkaloids. We repeated this analysis with root and petals from each of the three varieties, again with features that could be confidently assigned as iridoid, phenylpropanoid, or alkaloid (Supplementary Figs. 11-14). The analysis with root and petal protoplasts also showed the presence of a population of cells specializing in alkaloid accumulation, though specific patterns of flavonoid and secologanin accumulation varied among these tissues (Supplementary Figs. 11-14).

**Figure 3.**
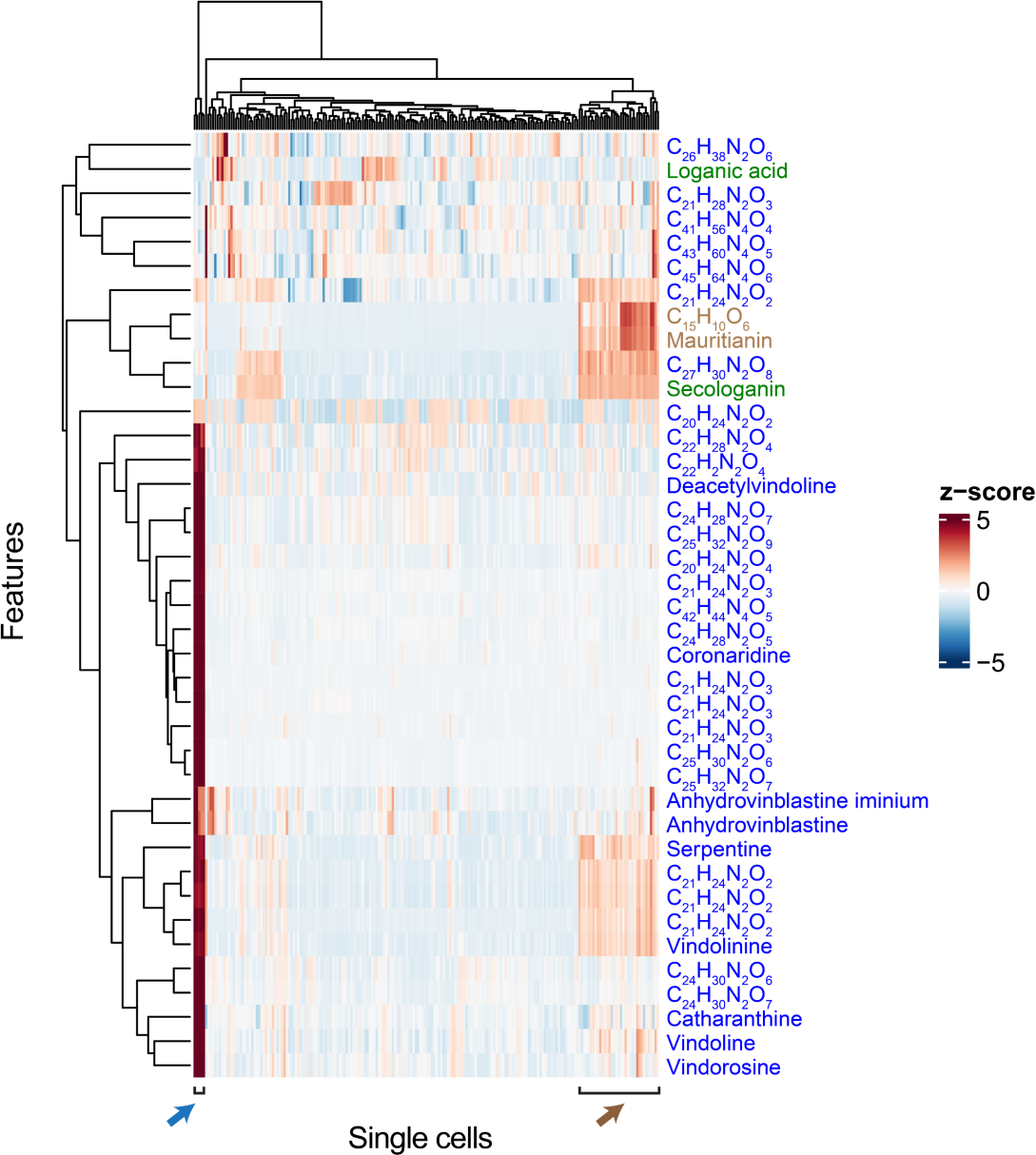
Hierarchical clustering analysis of 202 leaf protoplasts using a set of chemical features (39) that could be confidently assigned to iridoid (green compound names), alkaloid (blue compound names and formulas) or flavonoid (brown compound name and formula) types. The blue arrow indicates the group of cells with high levels of alkaloids. The brown arrow indicates the group of cells that accumulate primarily flavonoids and secologanin.

### Natural product concentrations across cell populations

This scMS workflow allowed simultaneous targeted and untargeted analysis of metabolites. Single cell analysis was performed in full scan mode, allowing sufficient scanning events for quantification, whilst a pooled quality control (QC) sample was used for fragmentation analysis to permit structural characterization. To accurately identify and quantify levels of metabolites in a single cell, we used external calibration curves of the authentic standards to convert the measured peak area for each compound into an absolute quantity. Additionally, the diameter of each cell was measured from the images acquired during the cell picking, allowing us to estimate the volume of the cell, which was then used to calculate the concentration of each of these molecules in an individual cell (Supplementary Data 4).

Strikingly, many cells contained millimolar concentrations of the compounds subjected to targeted monitoring. The iridoid monoterpene secologanin is by far the most abundant natural product observed in the bulk leaf tissue (12.5 mg g^−1^ fresh weight), and this is reflected by the fact that secologanin is found at high concentrations (50-600 mM) in a high percentage (ca. 30%) of leaf cells sampled (Fig. 4). Anhydrovinblastine, the precursor for vinblastine, is much less abundant in leaf, and this compound was observed at a range of 300 µM to 10 mM in a smaller number of cells (ca. 3% of all cells that were sampled). In contrast, vinblastine is present at low levels in bulk tissue, and correspondingly, was observed at a maximum of 100 µM concentration in only one leaf-derived cell out of all cells analyzed (Fig. 4, Supplementary Fig. 15). High variability in the levels of all quantified compounds was observed in these cell populations from all five tissues. For example, secologanin was detected at concentrations ranging from 5 mM to 60 mM in petal cells (Fig. 4). Notably, although a few leaf and root cells accumulated catharanthine to concentrations over 100 mM, the maximum concentration of catharanthine that can be reached in solution (pH 5, the expected pH of the vacuole) is ca. 20 mM (see Methods). Natural deep eutectic solvents have been proposed to aid in the solubilization of certain metabolites such as anthocyanins in plants^18–20^, and this may also be the case for alkaloids.

**Figure 4.**
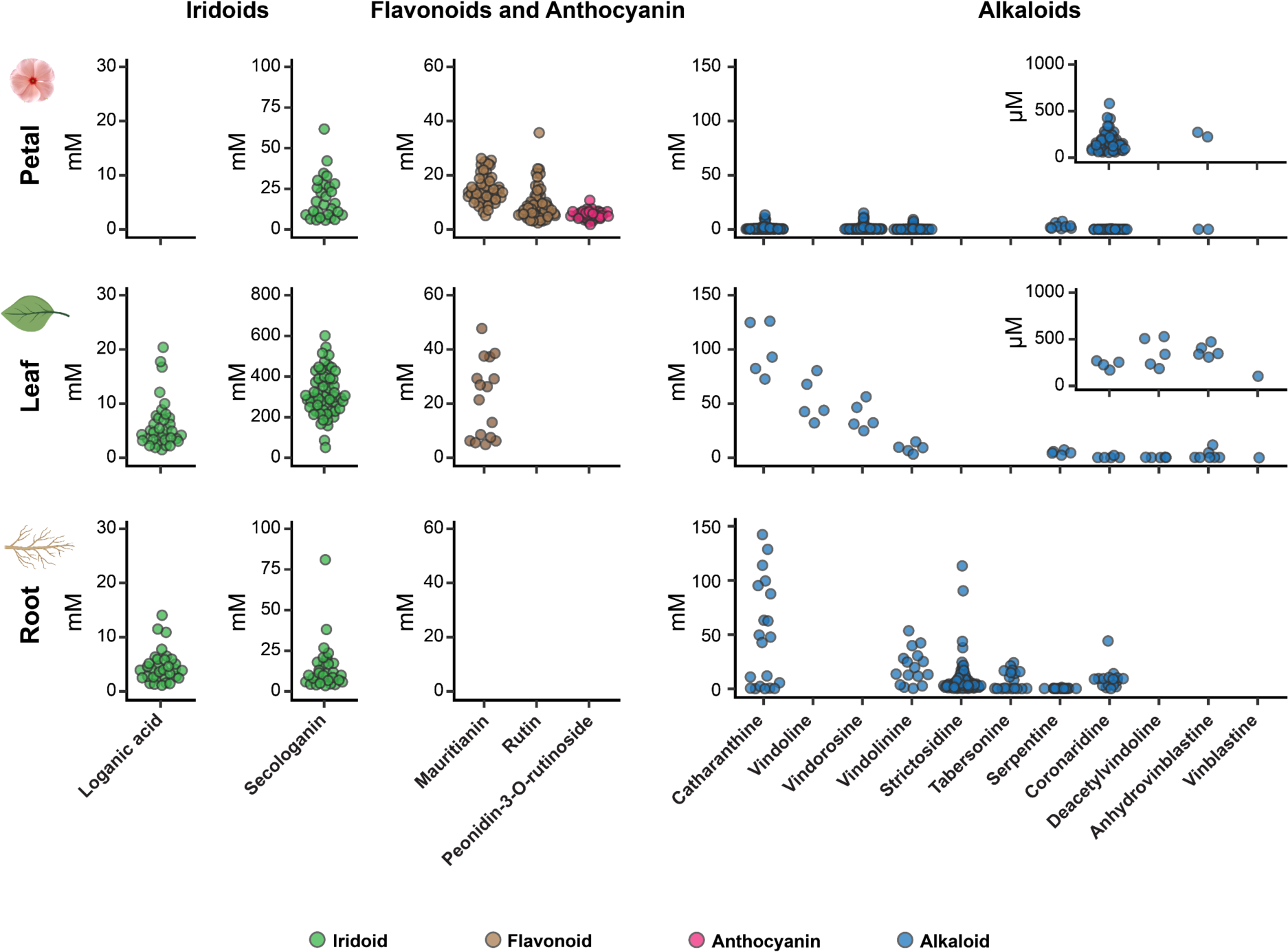
Quantification of 16 compounds for which standard curves of authentic standards could be generated. Plots show the concentration for each of these compounds in each cell. Colors of the data points represent the class of metabolites. All samples were from the Sunstorm Apricot (SA) cultivar.

### Cell type specificity compared among tissues

The untargeted metabolic analyses suggested the presence of sub-populations of cells that specialized in accumulating alkaloids or flavonoids (Fig. 3, Supplementary Figs. 11-14). To more accurately assess the ratio of compounds produced in each cell across the entire cell population that was measured, we used these quantitative data to generate stacked plots to show the absolute levels (mM) of each of the 16 quantified compounds in each cell (Fig. 5a, Supplementary Figs. 16b, 17, 20, 21). In leaf, we observed a sub-population of cells that specifically accumulate loganic acid (Fig. 5a). Since loganic acid has been shown to be synthesized in internal phloem-associated parenchyma (IPAP) cells^21^, we used the presence of this molecule as a marker for this cell type. Serpentine, an alkaloid that fluoresces under UV^5, 17^, was used as a marker for the assignment of idioblast cells, a rare cell type (2-3%^5^ of the total cell population), that accumulates the majority of the alkaloids. Therefore, the scMS data highlights that the high alkaloid levels observed in the bulk tissue are due to very few specialized cells containing large quantities of the compounds. Secologanin, which is observed in the bulk tissue at higher levels than any alkaloid, is detected in high concentrations in a much larger population of cells (Fig. 5a).

**Figure 5.**
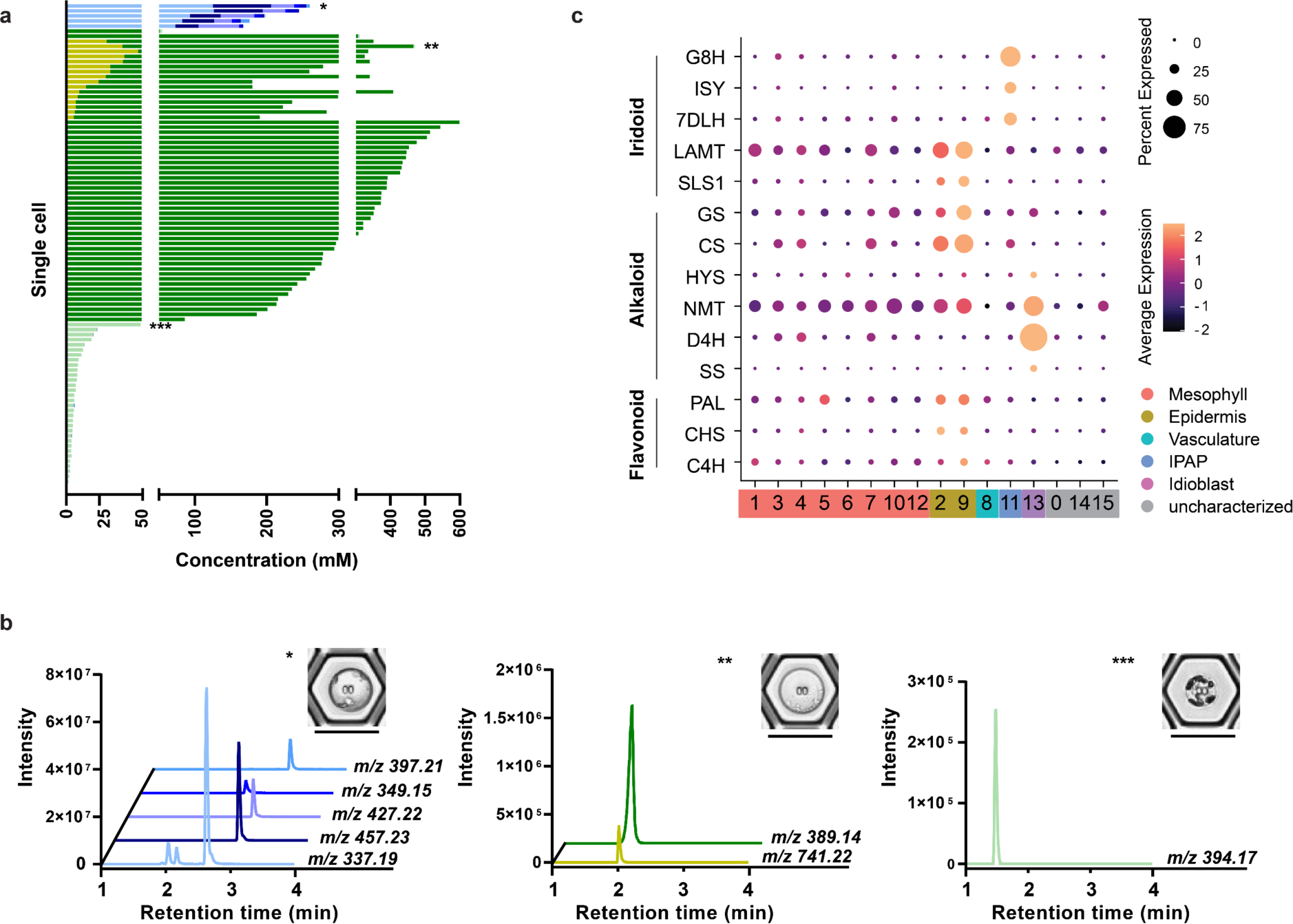
Ratio of compounds found across the population of leaf cells (202 cells). **a,** Stack plot showing the absolute concentration of each of the 16 quantified metabolites in each cell. Colors indicate classes of compounds: bars in blue shades represent alkaloids, dark green represents secologain (iridoid), light green represents loganic acid (iridoid) and yellow represents mauritanin (flavonoid). The asterisks indicate the cell for which the chromatograms in panel are shown in panel b. **b,** Representative chromatograms of these compounds from individual cells. (*m/z* 394.17, loganic acid; *m/z* 389.14, secologanin; *m/z* 741.22, mauritanin; *m/z* 337.19, vindolinine and catharanthine; *m/z* 457.23, vindoline; *m/z* 427.22, vindorosine; *m/z* 349.15, serpentine; *m/z* 397.21, anhydrovinblastine). The scale bar is 50 µm. **c,** scRNA-seq data (Sunstorm Apricot leaf) for selected biosynthetic genes in iridoid, alkaloid and flavonoid biosynthesis.

Notably, the partitioning of cells observed from the leaf-derived scMS data is only partially reflected in the corresponding scRNA-seq data (Fig. 5c). While the gene expression data shows that catharanthine is synthesized in epidermal cells, this alkaloid almost exclusively accumulates in cells assigned as idioblasts (*e.g.* catharanthine is found in the same cells as serpentine), suggesting that an efficient transport mechanism of catharanthine from epidermal to idioblast cells is in place. Additionally, the scMS data shows two distinct populations of cells that accumulate secologanin (green colored bars, Fig. 5a, b): one population that also accumulates the flavonoid mauritianin (yellow colored bars, Fig. 5a, b) and one that does not contain quantifiable amounts of any flavonoid-like compound (Fig. 5a). This distinction is not readily apparent in the scRNA-seq data, as phenylpropanoid biosynthetic genes^22^ (e.g. PAL, CHS, C4H) and secologanin biosynthesis genes (LAMT, SLS) were detected in the same cell clusters by scRNA-seq (Fig. 5c). Secologanin may serve a defensive role in addition to being a biosynthetic intermediate^23^; secologanin may therefore be transported to additional cell types after synthesis to support the additional biological function of this molecule^24^. On average, cells that accumulate both secologanin and flavonoid have lower levels of secologanin than cells that are specialized for secologanin (Fig. 5a).

We also compared scMS profiles between root and leaf tissue (Supplementary Fig. 16). As in leaves, roots have cell sub-populations that specialize in accumulating alkaloids (*e.g.* catharanthine). We also identified the root specific alkaloid hörhammercine that is derived from tabersonine (Fig. 1a); although this could not be quantified accurately due to scarcity of the authentic standard, we could definitively determine that this alkaloid co-localized with catharanthine (Supplementary Fig. 11), providing further support for the observation that alkaloids accumulate in specialized cell types in root analogous to leaves. A second cell sub-population accumulates iridoids and the upstream alkaloid strictosidine, while a third sub-population of cells accumulates only strictosidine, but no iridoids. Flavonoids were not detected in root-derived cells. These three distinct populations are not apparent from the scRNA-seq data, which shows that iridoid and alkaloid biosynthetic genes are expressed in ground cells (Supplementary Fig. 16a). Therefore, the root scMS data reveal cell type specificity that is not detectable from the scRNA-seq data. The mechanism by which the pattern of metabolite accumulation observed in the scMS dataset is established remains to be determined.

Finally, we examined flower petals, which contain flavonoids, anthocyanins, iridoids, and alkaloids. In cells derived from petals of the SA cultivar, the majority of cells sampled are highly enriched in either rutin (flavonoid)/peonidin 3-*O*-rutinoside (anthocyanin), the flavonoid mauritianin or a combination of monoterpene indole alkaloids (Supplementary Fig. 17). We also observe a fourth, smaller subpopulation of cells that are specialized in secologanin accumulation. To compare the scMS data with gene expression profiles, we also generated a scRNA-seq dataset for petals (Supplementary Fig. 18, Supplementary Data 5). Surprisingly, many alkaloid and iridoid biosynthetic genes are expressed at negligible levels in flower petals (Supplementary Fig. 18). This was consistent with bulk RNA-seq data taken at time points after flower opening (Supplementary Fig. 19, Supplementary Data 6). Petal provides a striking case in which the prevalence of the metabolites – which are found in high levels in this tissue– is not correlated with gene expression. The metabolites, or biosynthetic intermediates of these metabolites, that are detected in petal-derived cells may be synthesized during different stages of flower development, or alternatively, these compounds could be transported from other tissues.

We also investigated petal-derived cells of LBE and ABH cultivars by scMS (Supplementary Fig. 20 and 21). In both LBE and ABH, cells that are specialized to accumulate alkaloids are observed. However, while SA and ABH have cells that specialize in secologanin accumulation, in LBE, secologanin nearly always co-localizes with flavonoids. Of all three cultivars, LBE has the highest levels of alkaloids in petal, and this is reflected in the single cell data, with concentrations of 300-400 mM being reached for the total alkaloid level (Supplementary Fig. 20). The mechanisms by which these localization patterns are achieved, or whether these different metabolite co-localization patterns have functional or ecological significance, remains to determined. Nevertheless, this scMS analysis clearly shows that leaf, root and petals of three cultivars store metabolites differently at the cell-type level.

To more easily visualize the differences in metabolite cell-type specificity across these five samples, we grouped the cell populations of each tissue into four clusters (using k-means clustering analysis) based on the peak area of the 20 natural products that could be structurally assigned with high confidence (Fig. 6, Supplementary Fig. 22). While these 20 compounds represent only a small fraction of the natural product profile diversity of *C. roseus*, this analysis shows that all tissues have cells that specialize in monoterpene indole alkaloid accumulation. However, iridoid, flavonoid, and anthocyanin compounds show different co-localization patterns across these tissues.

**Figure 6.**
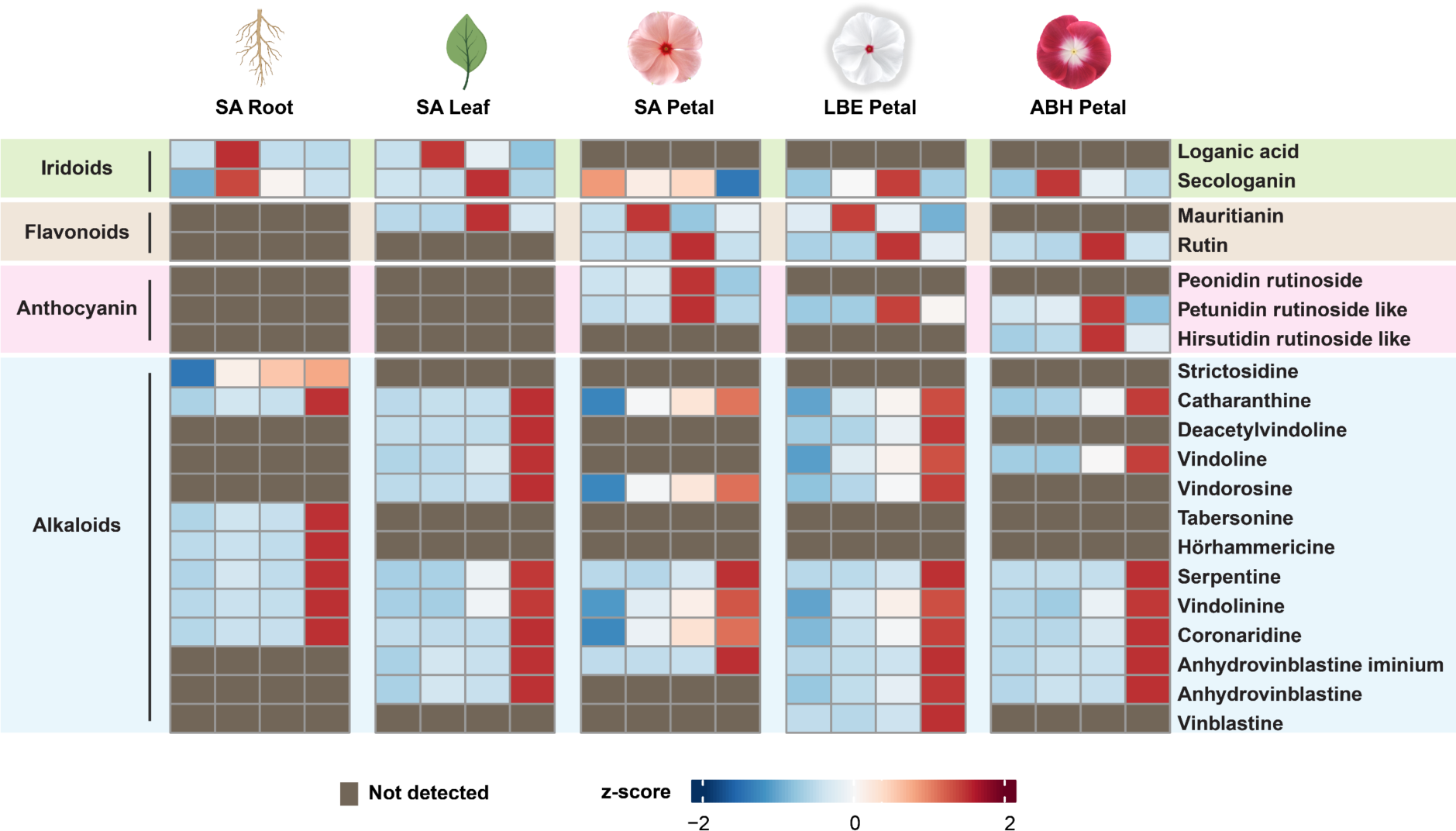
Comparison of metabolite profiles of cell populations across tissues and cultivars. Cells were grouped into four clusters based on the levels of the 20 metabolites that could be confidently assigned using k-means clustering analysis (Supplementary Fig. 22). The intensity of the compounds represented in the heat map is obtained by measuring, log-transforming, normalizing and averaging peak area of all cells in one cluster. The identity of all compounds except two were validated with an authentic standard. Authentic standards are not available for petunidin rutinoside-like and hirsutidin rutinoside-like compounds.

## Discussion

The medicinal plant *C. roseus* produces a wide array of natural products, including iridoid-type monoterpenes, flavonoids, anthocyanins, and monoterpene indole alkaloids^25, 26^. The importance of two of these alkaloids, anhydrovinblastine and vinblastine, which are used in cancer treatment^27^ have made *C. roseus* one of the most studied medicinal plants. Discovery and engineering of the biosynthetic pathways responsible for construction of complex natural products such as vinblastine is of paramount importance to improve access to these valuable plant-derived molecules (*e.g.* Zhang, J. *et al*.^28^).

The availability of scRNA-seq datasets, which reveal how biosynthetic elements are expressed in specific cell types, is transforming how we elucidate and engineer plant-derived natural product pathways. However, since the ultimate cellular location of natural products is not always accurately predicted by scRNA-seq, these transcriptomic datasets only provide a partial snapshot of natural product biosynthesis. A complete picture requires comparison of metabolite and biosynthetic gene localization. Here, we report a method for single cell mass spectrometry-based metabolomics (scMS) that can detect and quantify the iridoid-type monoterpenes, flavonoids, anthocyanins, and monoterpene indole alkaloids in individual cells from three different tissues (leaf, root and petal) of *C. roseu*s. Although we use a Q-Exactive Plus Orbitrap mass spectrometer in the work presented here, this scMS workflow is compatible with a variety of mass spectrometry instruments, making this method accessible for many laboratories. This approach is also, in principle, applicable for detection of any metabolite that can be detected using UHPLC-MS.

We performed both targeted and untargeted scMS analyses for leaf, root and petal cells, and then compared each of these scMS datasets to the corresponding scRNA-seq data from the same tissue. The comparison of the scMS and scRNA-seq datasets in each tissue highlighted distinct differences between metabolite location and biosynthetic gene expression. In leaf, the scMS data showed three major populations of cells, largely correlating with the scRNA-seq data. However, we saw that some alkaloids (*e.g.* catharanthine) were transported to a different cell type. Additionally, we noticed two distinct populations of cells that contained secologanin that was not immediately apparent from the biosynthetic gene expression profiles observed in the scRNA-seq data. Additionally, comparison of scMS and scRNA-seq data of root and petal suggested that in these tissues, expression of biosynthetic genes do not always correlate with the site of metabolite accumulation. The expression of biosynthetic genes in root suggested that iridoids and downstream alkaloids would be found in the same cell type (ground cells), but the scMS data showed that alkaloids are not found in the same cell type as the upstream iridoids. Although high levels of alkaloids were observed in petal cells, the biosynthetic gene expression was low, suggesting that these compounds are transported from other tissues, or that synthesis occurs at different stages of flower development. Overall, the comparison of scRNA-seq and scMS data strongly suggest the presence of many active intercellular transport processes. To date, only one inter-cellular transporter has been identified in *C. roseus*, the transporter responsible for movement of loganic acid from IPAP cells to epidermal cells^24^. This scMS and scRNA-seq data provides the foundation for the discovery of additional transporters.

The targeted scMS data show that individual cells can accumulate exceptionally high (> 100 mM) concentrations of many of these natural products. Glucosinolates may also reach millimolar levels in cells^29, 30^, but this is indirectly inferred from bulk tissue measurements. Most of the metabolites that accumulate to these high concentrations, such as secologanin and vindoline, are predicted to be localized in the vacuole^31, 32^. The high levels of loganic acid may also be because transport from IPAP to epidermal cells is a rate determining step. Iridoids, monoterpene indole alkaloids and phenylpropanoids all have unstable biosynthetic intermediates in the pathway, but none of these were observed in these scMS experiments. ScMS with protoplasts fed with isotopically labeled precursors may facilitate the identification of lower abundance compounds in these experiments.

Both targeted and untargeted MS suggest that the majority of monoterpene indole alkaloids accumulate in specialized cells that represent only a small fraction of the total cell population. The iridoid secologanin is found in high concentrations across a larger fraction of sampled leaf cells, highlighting the capacity of the plant cell factory to produce and store exceptionally high levels of a range of complex natural products. The most medicinally valuable alkaloid, vinblastine, is found at micromolar concentrations in only a small fraction of cells that were sampled, which reflects the low levels observed in the bulk tissue.

Although single cell sequencing is now being used broadly, rigorous structural characterization and quantification of metabolites in single cells has proven to be challenging. Here we report a robust method for single cell mass spectrometry in which we identify and quantify the concentrations of 16 metabolites across four natural product classes in individual cells of leaf, root, and petal of the medicinal plant *C. roseus*. In combination with scRNA-seq data, this scMS approach allows us to dissect the logistics of these pathways at a highly resolved level. Incorporation of this robust scMS pipeline into a single cell omics analysis pipeline can be a step change in how we understand the biosynthesis and biological function of these molecules.

## Acknowledgements

This work was supported by funds from the Max Planck Gesellschaft (S.E.O., L.C.), the Leibniz Prize, Deutsche Forschungsgemeinschaft (DFG, German Research Foundation) – 505457618 (S.E.O.), Georgia Research Alliance (C.R.B.), Georgia Seed Development (C.R.B.), University of Georgia (C.R.B.), and National Science Foundation (MCB-2309665 to C.R.B. and C.L.). We thank Brieanne Vaillancourt for assistance in sequence deposition to NCBI. We thank Omar Kamileen and Abdullah Sandhu for providing some authentic standards for the study. We would like to thank Dr. Ling Chuang and Dr. Marine Vallet for advices about data analysis and data visualization. Part of the workflow in Fig. 2 and plant art were created with BioRender.com.

## Methods

### Chemicals

All solvents used in this study were of UPLC/MS grade. Information about chemicals and reagents is listed in Supplementary Table 6.

### Plant growth conditions

*Catharanthus roseus* (*C. roseus*) plants of Sunstorm Apricot (SA), Little Bright Eyes (LBE) and Atlantis Burgundy Halo (ABH) cultivars were germinated and grown in a York chamber at 23 °C, under a 16h/8h light/dark cycle.

### Protoplast isolation

#### Leaf protoplast isolation

A healthy plant was watered and left in the dark the day before the leaves were harvested. Three leaves of ca. 3 cm in length (Fig. 1b) were selected, rinsed gently with water, and cut in 1 mm strips with a sterile surgical blade. The leaf strips were immediately transferred to a Petri dish with 10 mL of digestion medium (2% (w/v) Cellulase Onozuka R-10, 0.3% (w/v) macerozyme R-10 and 0.1% (v/v) pectinase dissolved in Mannitol-MES (MM) buffer. MM buffer contained 0.4 M mannitol and 20 mM MES, pH 5.7-5.8, adjusted with 1 M KOH. The open Petri dish was put inside a desiccator and 100 mBar vacuum was applied for 15 min to infiltrate the medium into the leaf strips. The vacuum was gently disrupted for 10 s after every 1 min. The leaf strips were then incubated in the digestion medium for 2.5 h at room temperature. After the incubation, the Petri dish was placed on an orbital shaker at ca. 70 rpm, for 30 min at room temperature to help release the protoplasts. The protoplast suspension was filtered through nylon sieves of 70 μm and then 40 μm to remove cellular debris. After that, the suspension was transferred to two 15 mL conical tubes. The protoplast suspension was centrifuged at 70 x g with gentle acceleration/deceleration, for 5 min, at 23 °C to pellet the protoplasts. The supernatant was removed and the protoplasts were washed three times with 5 mL of MM buffer. Finally, the protoplasts were pooled together and resuspended in 1 mL of MM buffer. The protoplast concentration and viability were determined using a haemocytometer and fluorescein diacetate staining, respectively. The final concentration of protoplasts was adjusted to 10^6^ protoplasts in 1 mL.

#### Petal protoplast isolation

The petals were removed from fully opened flowers (stage 1, Supplementary Fig. 19) and cut into 1 mm strips with a sterile surgical blade. Protoplast isolation was performed using the same protocol used for the leaves, except for the time of incubation in the digestion medium, which was decreased to 1.5 hours.

#### Root protoplast isolation

Roots from a young healthy plant (6-7 weeks old) were used for protoplasting (Fig. 1b). After removing the soil, the roots were washed with water and gently dried to avoid damaging the tissue. The roots were then finely sliced with a sterile surgical blade. Protoplast isolation was performed using the same protocol used for the leaves, with a few modifications: the concentration of macerozyme in the digestion solution, infiltration time, incubation time and centrifugation speed were optimized to 0.6%, 30 minutes, 1.5 hours and 200 g, respectively, for root protoplasting.

### Single cell picking

SIEVEWELL™ chips (Sartorius) with 90,000 nanowells (50 μm x 50 μm, depth x diameter) were used for single-cell trapping and sorting. Chips were primed with 100% ethanol and washed twice with MM buffer. The chips were then incubated with 5% BSA in MM buffer for 30 minutes at room temperature. Subsequently, the 5% BSA in MM buffer solution was discarded through the side port and replaced with MM Buffer. Finally, 1 mL of diluted protoplasts suspension (approximately 10,000 cells) was carefully added and dispensed in a Z-shape across the chip. Liquid (1 mL) was discarded from the side ports.

The SIEVEWELL™ chip was then mounted on the CellCelector™ Flex (Sartorius) instrument and the cells were visualized using the optical unit, constituted by a fluorescence microscope (Spectra X Lumencor) and a CCD camera (XM-10). Photos in bright-field or fluorescence (DAPI filter) were acquired, depending on the experiment. Single protoplasts were picked together with 20 nL of well solution using a 50 μm glass capillary and dispensed into SureSTART™ WebSeal™ 96-Well Microtiter plates (Thermo Fisher Scientific) containing 6 μL of MilliQ water. Pictures of the nanowells before (containing a single cell) and after picking (without a cell) were recorded. After picking, 6 μL of MeOH containing 20 nM ajmaline (internal standard) was added to each well. Pooled QC samples consisting of 2 µL of each sample were made for each experiment and used for quality control and for MS^2^ (fragmentation) analysis.

### Preparation of bulk tissue extracts

We performed untargeted and targeted mass spectrometry analysis of bulk tissue extracts of leaves, roots and petals of *C. roseus* SA, a cultivar with peach petals (Fig. 1b). Bulk tissue samples were diluted so that the concentration of the alkaloid catharanthine matched the concentration ranges observed in the single cell datasets. We also characterized the metabolic profiles of petals from LBE and ABH cultivars, with white- and burgundy-colored petals, respectively (Fig. 1b). All tissues were ground to a fine powder using a Tissuelyser II (Qiagen). Metabolites were extracted with pure MeOH containing 2 µM ajmaline as an internal standard at a ratio of 300 µL of solvent per 10 mg of tissue. After vortexing and sonication for 10 min, the tissue extracts were filtered through a 0.2 μm PTFE filter. The extracts from leaf, root and petal were diluted 500-fold, 200-fold and 50-fold, respectively before the analysis. Pooled QC samples of the extracts were also prepared.

### LC-MS analysis

UHPLC-HRMS analysis was performed on a Vanquish (Thermo Fisher Scientific) system coupled to a Q-Exactive Plus Orbitrap (Thermo Fisher Scientific) mass spectrometer. For metabolite separation, a Waters™ ACQUITY UPLC BEH C18 130 Å column (1.7 µm, 1 mm x 50 mm) was used at a temperature of 40 °C. The binary mobile phases were 0.1% HCOOH (formic acid) in MilliQ water (aqueous phase) (A) and acetonitrile (ACN) (B). The gradient elution started with 1% ACN and increased linearly to 70% ACN over 5 minutes. The wash stage was performed at 99% ACN for 0.5 minutes before switching back to 1% ACN for 1.5 minutes to condition the column for the next injection. Total time for chromatographic separation was 7 minutes. The flow rate was 0.3 mL min^−1^ during the chromatographic separation. Injection volume was 4 μL. The autosampler was kept at 10 °C throughout the analysis. The needle in the autosampler was washed using a mixture of methanol and MilliQ water (1:1, v:v) for 20 seconds after draw and at a speed of 50 µL s^−1^.

The Q-Exactive Plus Orbitrap mass spectrometer (Thermo Fisher Scientific) was equipped with a heated electrospray ionization (HESI) source. The mass spectrometer was calibrated using the Pierce positive and negative ion mass calibration solution (Thermo Fisher Scientific). The operating parameters of HESI were based on the UHPLC flow rate of 0.3 mL min^−1^ using source auto default: sheath gas flow rate at 48; auxiliary gas flow rate at 11; sweep gas flow rate at 2; spray voltage +3,500 V; capillary temperature at 256 °C; auxiliary gas heater temperature at 413 °C; and S-lens RF level at 50. Acquisition was performed in full scan MS mode (resolution 70,000-FWHM at 200 Da) in positive mode over the mass range *m/z* from 120 to 1,000. The full-scan and data-dependent MS/MS mode (full MS/dd-MS^2^ Top10) was used for QC pooled samples to simultaneously record the spectra of the precursors as well as their MS/MS (fragmentation). Besides, the full MS/dd-MS^2^ mode with inclusion list was also applied for the pooled QC samples to confirm fragments of the selected precursors. The parameters for dd-MS^2^ were set up as follows: resolution 17,500, mass isolation window 0.7 Da and normalized collision energy (NCE) was set at 3 levels: 15%, 30% and 45%. Spectrum data format was centroid. All the parameters of the UHPLC-HRMS system were controlled through Xcalibur software version 4.3.73.11 (Thermo Fisher Scientific).

### Method partial validation and quantification

The repeatability of the method was tested by evaluating the stability of the retention time and detected peak areas of selected compounds during ten injections of the bulk methanolic extracts of three different tissues for three consecutive days. The inter-day precision was performed by applying the intra-day injections for three consecutive days (Supplementary Table 2).

Alkaloids, iridoids, flavonoids and anthocyanins standard solutions were prepared in MeOH at a concentration of approximately 1 mM (exact concentration was recorded). Serial dilutions were made until 0.001 nM and analyzed by UHPLC-MS to determine limit of quantification (LOQ) and calibration range (Supplementary Table 4). Each calibration point was measured in triplicate. The extracted peak areas were used to calculate linear regression curves. Chromatographic peak area from extracted ion chromatograms (EIC) were integrated and extracted using the Xcalibur Quan Browser version 4.3.73.11 (Thermo Fisher Scientific).

### LC-MS data processing and analyzing

Raw data were imported into Compound Discoverer™ software 3.2 (Thermo Fisher Scientific) for peak picking, deconvolution and formula assignment. Parameters are listed in Supplementary Table 7.

The identity of selected ions was confirmed by comparison of their retention time and fragmentation spectra with analytical standards. Assignment of putative identities was performed with other glucosides with the same aglycon for two additional anthocyanin compounds. All the relevant features with MS^2^ data were further analyzed with SIRIUS 5 software^11^. CANOPUS^12^ was used to predict compound classes from mass spectra. Chemical classification of all features was performed using NPclassifier^13^ results and only classification with natural product pathway probabilities of > 0.8 was selected.

### Catharanthine solubility experiment

14.07 mg of catharanthine powder was dissolved in 20 µL of DMSO. After all the powder was dissolved, the catharanthine solution was added in 0.1 µL additions to 100 µL acetate buffer 0.1 M, pH 5. The maximum volume of catharanthine solution that could be dissolved in the buffer was 1 µL. The final concentration of dissolved catharanthine was approximately 21 mM.

### Bioinformatics analysis

The feature list from Compound Discoverer™ was exported in .xlsx format. The peak areas were log-transformed and then scaled before performing principal components analysis (PCA) for reducing dimension, hierarchical clustering and k-means clustering analysis^33^. Statistical analyses were performed using R (version 4.3.1). The heatmaps were visualized by pheatmap package (version 1.0.12) and ComplexHeatmap package (version 2.18.0)^34^. Violin point graphs were visualized by the ggplot2 (version 3.5.0) package. Stacked bar charts were visualized by GraphPad Prism software (version 9.5.1).

### Bulk RNA sequencing of petals

Petals were collected from flowers at three different stages of development (Supplementary Fig. 19) and flash frozen in liquid nitrogen. Three biological replicates were prepared for each stage. After grinding with a Tissuelyser II (Qiagen), the total RNA was extracted using the RNeasy Plant kit (Qiagen) and sequenced by Biomarkers Technologies GmbH on an Illumina Novoseq X (PE150) platform.

### Single Cell RNA-sequencing

#### Leaf and Root

For assessing single-cell transcriptome of leaf and root tissues, we reanalyzed 10x Genomics datasets from the previous study of *C. roseus*^5^ which is publicly available at https://doi.org/10.5061/dryad.d2547d851. Downstream analysis and visualization were performed as described previously.

#### Petal

Protoplasts from *C. roseus* SA petals were isolated as described above, using four flowers for each biological replicate. ScRNA-Seq libraries were constructed using the PIPseq v4.0 Plus kit (Fluent Biosciences) according to the manufacturer’s instructions, targeting ∼ 3,154 (cro_bz) and ∼ 3,306 cells (cro_ca)^35^ The libraries were sequenced on an Element Biosciences Aviti Instrument. For processing reads, pipseeker v3.1.3 (Fluent Biosciences) barcode pipeline was adopted to trim read 2 and detect barcode whitelist. Reads were then aligned to *C. roseus* (v3.0^5^) genome using the STARsolo pipeline (v2.7.10) with following parameters^36^: -- soloBarcodeReadLength 0 --alignIntronMax 5000 --soloUMIlen 12 –soloCellFilter EmptyDrops_CR --soloFeatures GeneFull --soloMultiMappers EM --soloType CB_UMI_Simple --soloCBwhitelist using the barcode whitelists detected by pipseeker. Ambient RNA was removed by DecontX^37^, and cells harboring RNA between 300 and 6,000 were kept for downstream analyses in seurat v5.0.1^38^. After log-normalization, top 3,000 variable genes were selected for integrating two replicates. Uniform manifold approximation and projection (UMAP) were calculated using the first 30 principal components and the same clustering parameters as leaf and root dataset. Cell types were annotated based on marker genes (vasculature, epidermis, idioblast) of previous studies^5, 39^, and functional annotations of *de novo* marker genes (epidermis, parenchyma) in flower dataset.

### Microscopy

Plant organ micrographs were acquired as tile scans (2×2 or 3×3) with a AXIO Zoom.V16 (Zeiss) equipped with a PlanApo Z 0.5x objective and a custom light box for homogenous indirect illumination. The tile scans were aligned and fused in ZEN (Zen lite 3.4, Zeiss) and background correction was done in Photoshop (Creative Cloud 2024, Adobe Inc.) after exporting.

Plant organ sections (approximately 50 µm thick) were prepared with a manual rotation Plant microtome (NK System MTH-1) and micrographs were subsequently acquired using an Apo Z 1.5x objective (Zeiss) with transmitted light illumination at the AXIO Zoom.V16. Micrographs of the protoplasts were acquired at an Imager.Z1 (Zeiss) equipped with a 20x/0.8 Plan-APOCHROMAT objective and DIC illumination. Lambda scans of the protoplasts were acquired with a cLSM 880 (Zeiss) equipped with a 20x/0.8 Plan-APOCHROMAT objective and 405nm laser diode (10% transmission) or 633 nm Helium-Neon laser (50% transmission) for excitation. Emission spectra were detected in lambda mode with a 9 nm binning ranging from 414 nm to 655 nm and 750 nm detector gain. The displayed micrographs are coded to show the actual color of the maximum intensity signal for each pixel.

## Supplementary Information

### Supplementary Figures

**Supplementary Figure 1.**
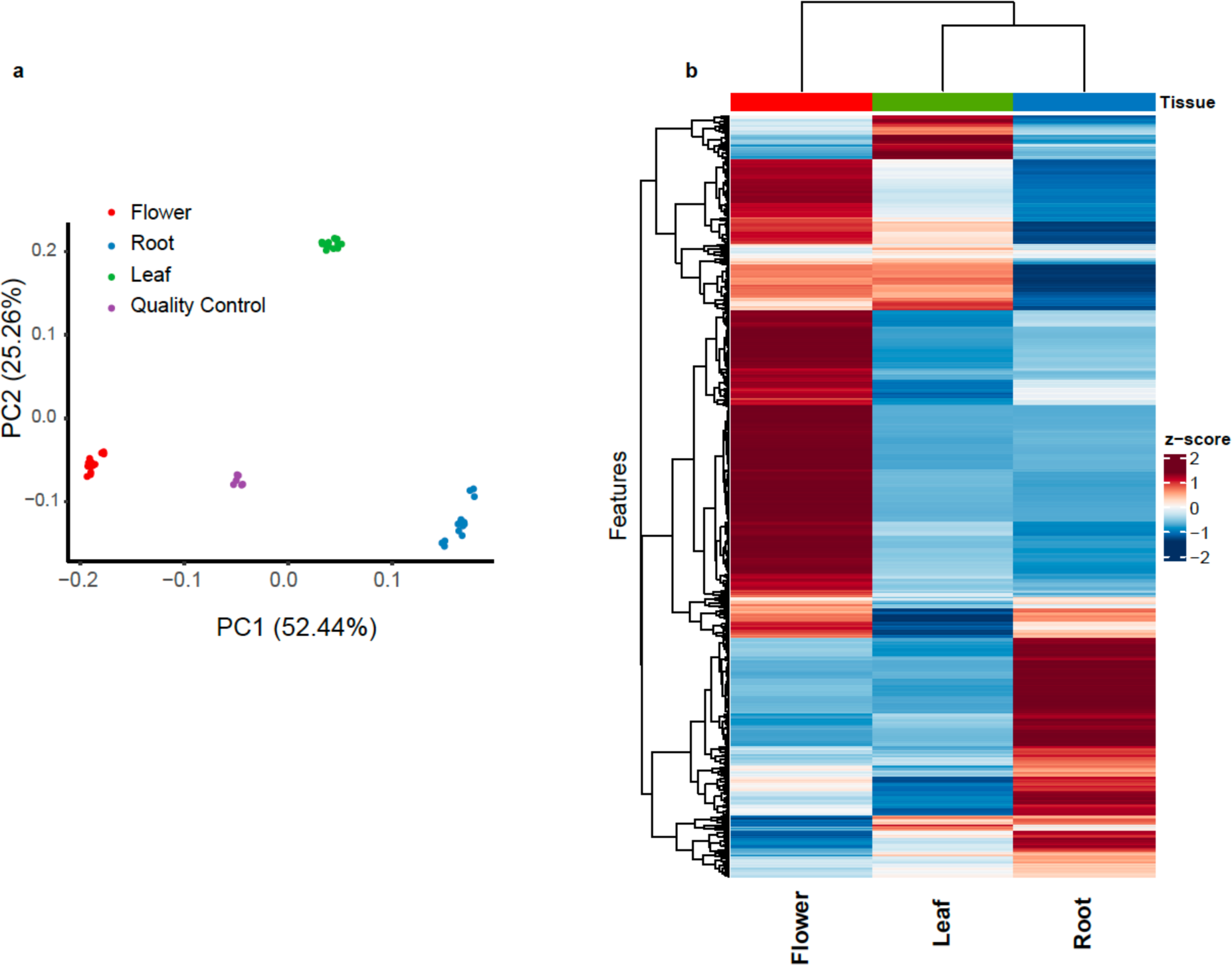
Metabolomic analysis of *C. roseus* SA leaf, root and petal bulk tissue extracts. **a**, PCA plot of the chemical features (formula assignment) shows that the chemical composition varies substantially among the three tissues. **b**, Heatmap showing the occurrence of the different features across the three tissues. 5 biological replicates, 3 technical replicates for each biological replicate and 1014 features are listed (Supplementary Data 1).

**Supplementary Figure 2.**
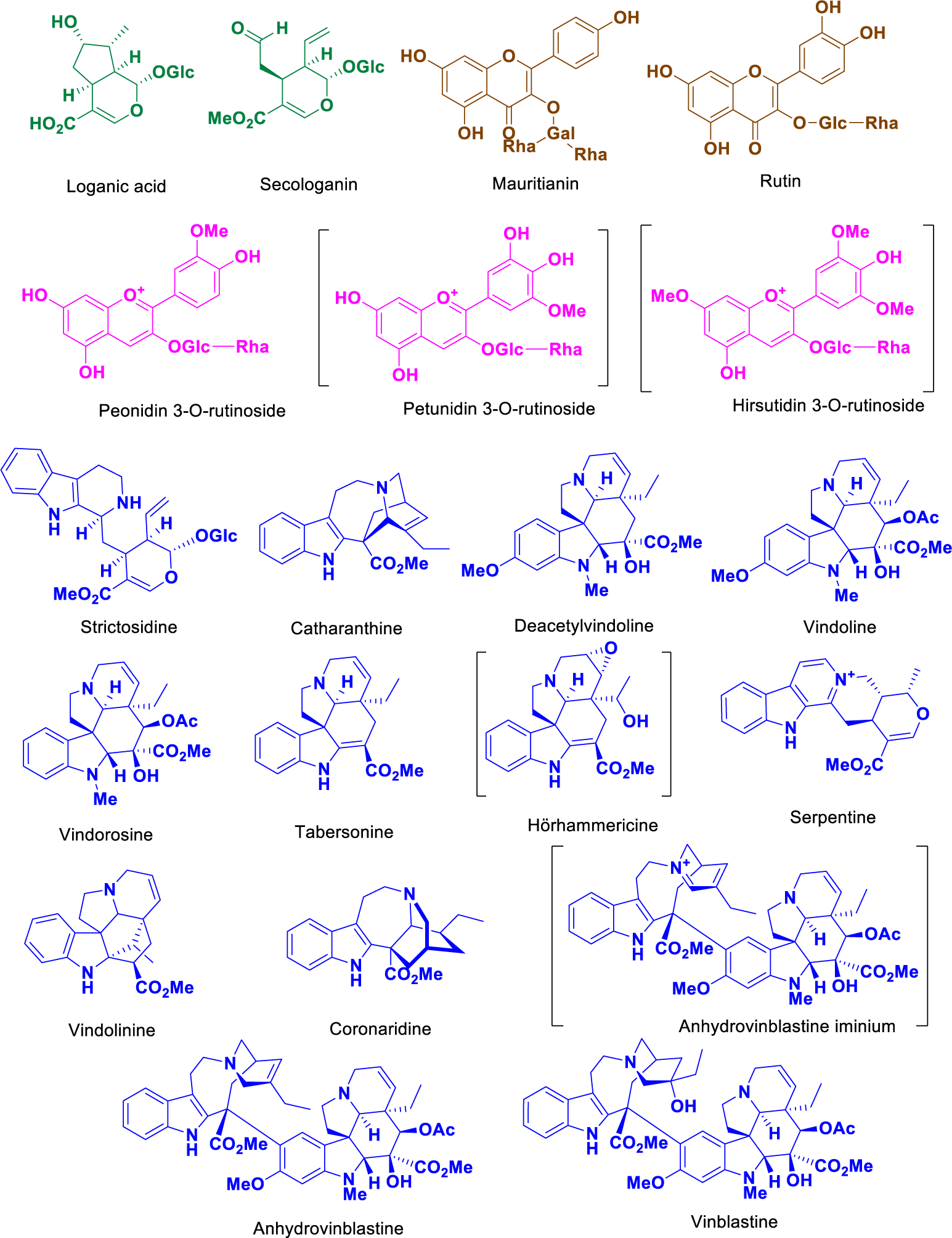
Chemical structures of the compounds identified in this study. The compounds in brackets were not quantified. Petunidin-3-O-rutinoside and hirsutidin-3-O-rutinoside are structural assignments based on mass and fragmentation. Authentic standards for anhydrovinblastine iminium and hörhammercine are available, but not in quantities sufficient for external calibration curves needed for quantification. All compounds not in brackets were quantified in the scMS studies using external calibration curves of authentic standards.

**Supplementary Figure 3.**
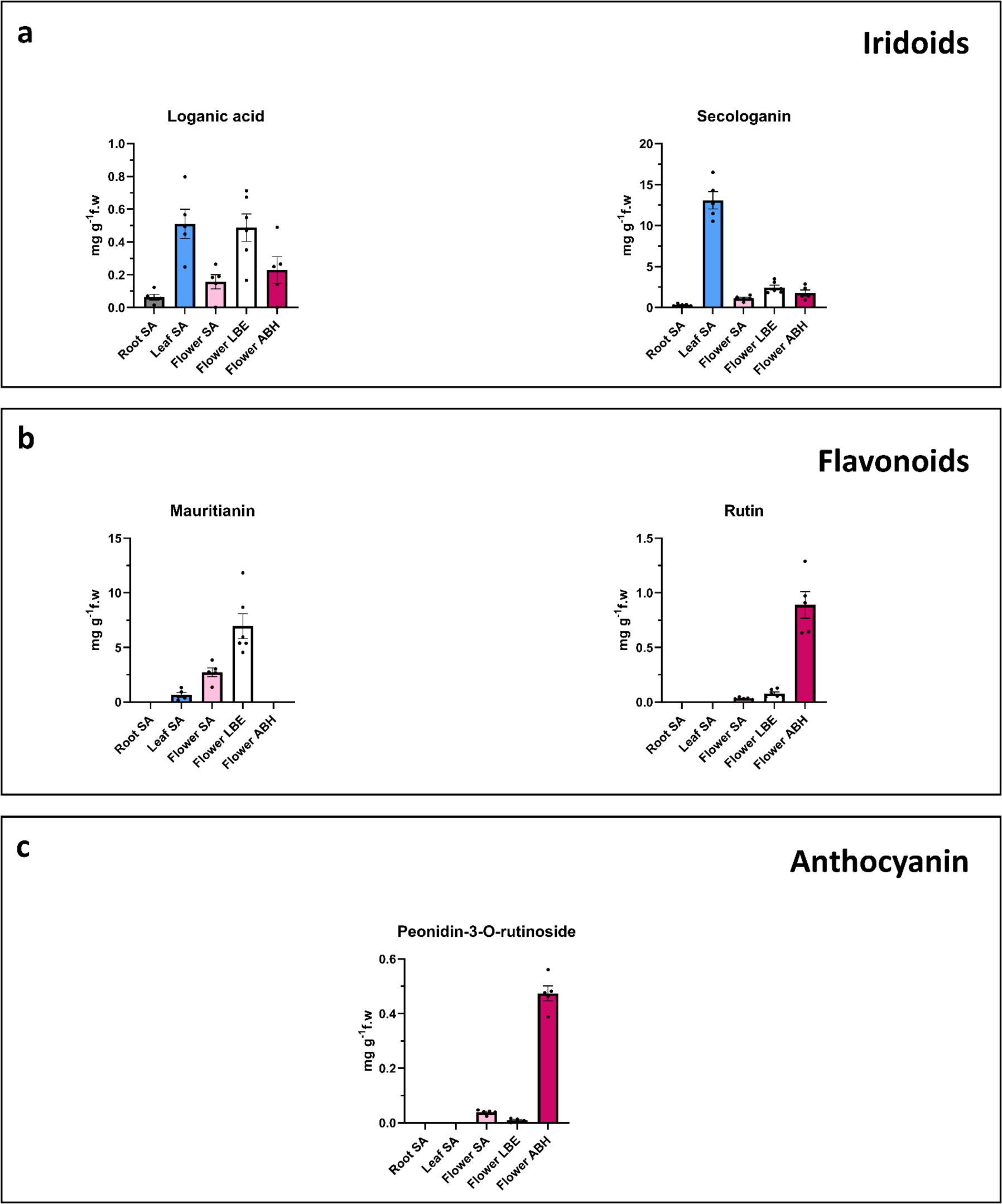
Quantification of key **a**, iridoids, **b**, flavonoids and **c**, the anthocyanin peonidin-3-O-rutinoside in bulk tissues.

**Supplementary Figure 4.**
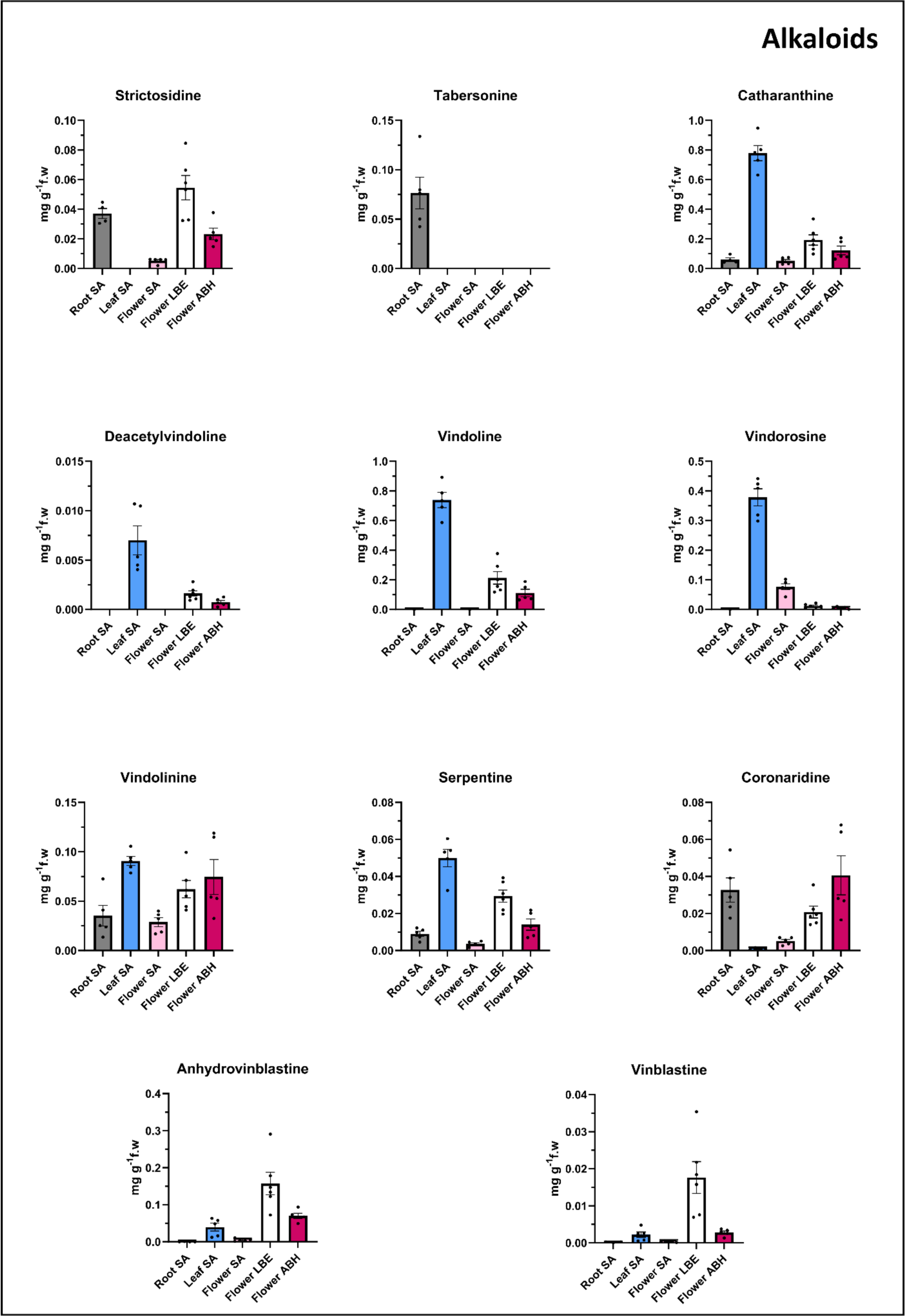
Quantification of alkaloids in bulk tissues.

**Supplementary Figure 5.**
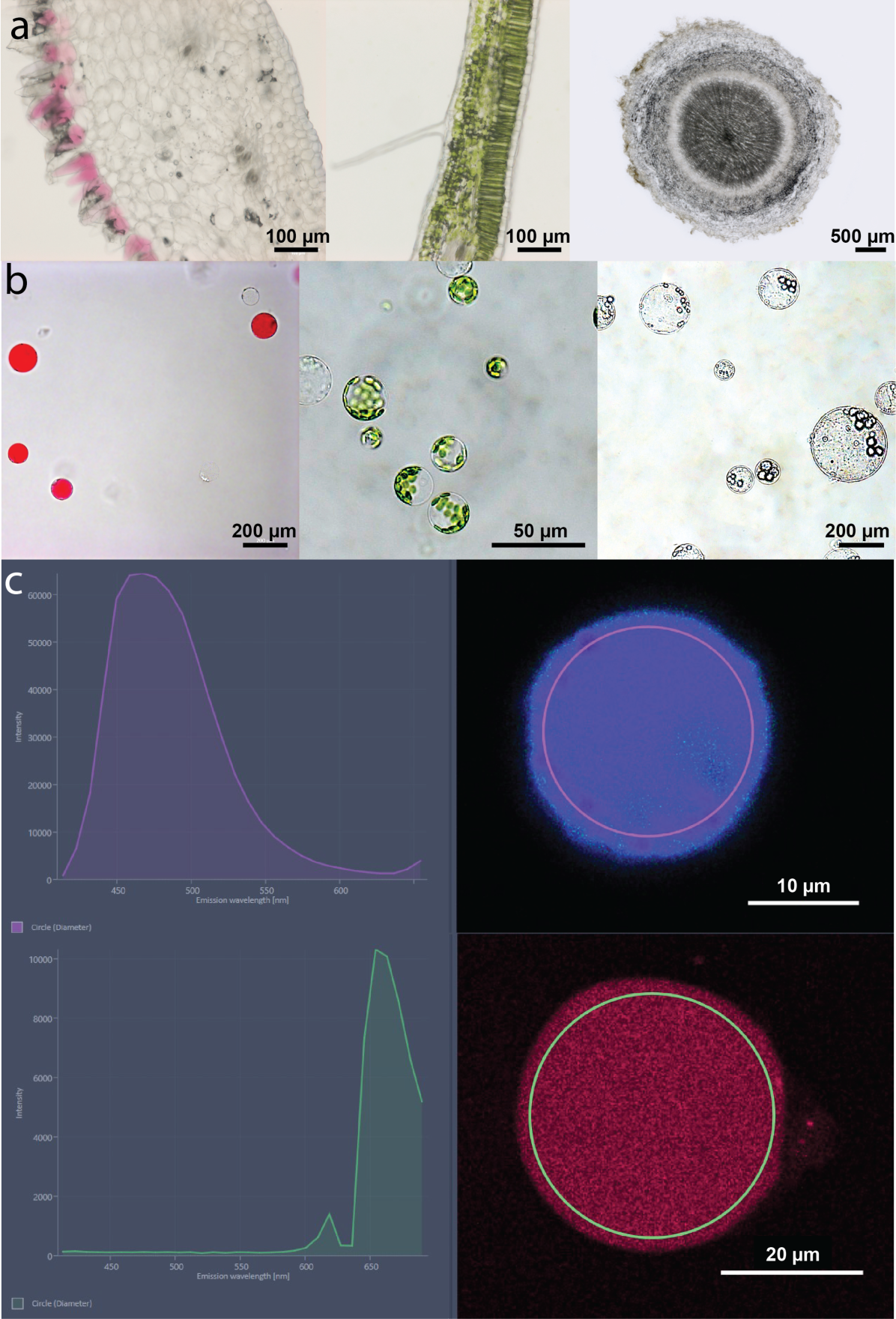
**a,** Microscopic photos of representative tissue sections (from left to right): SA petal, SA leaf, SA root. **b**, Photos of representative captured protoplasts (from left to right): ABH petal protoplasts, SA leaf protoplasts, SA root protoplasts. **c,** Photos of representative idioblast (top) and pigment (bottom) cells with their emission spectra.

**Supplementary Figure 6.**
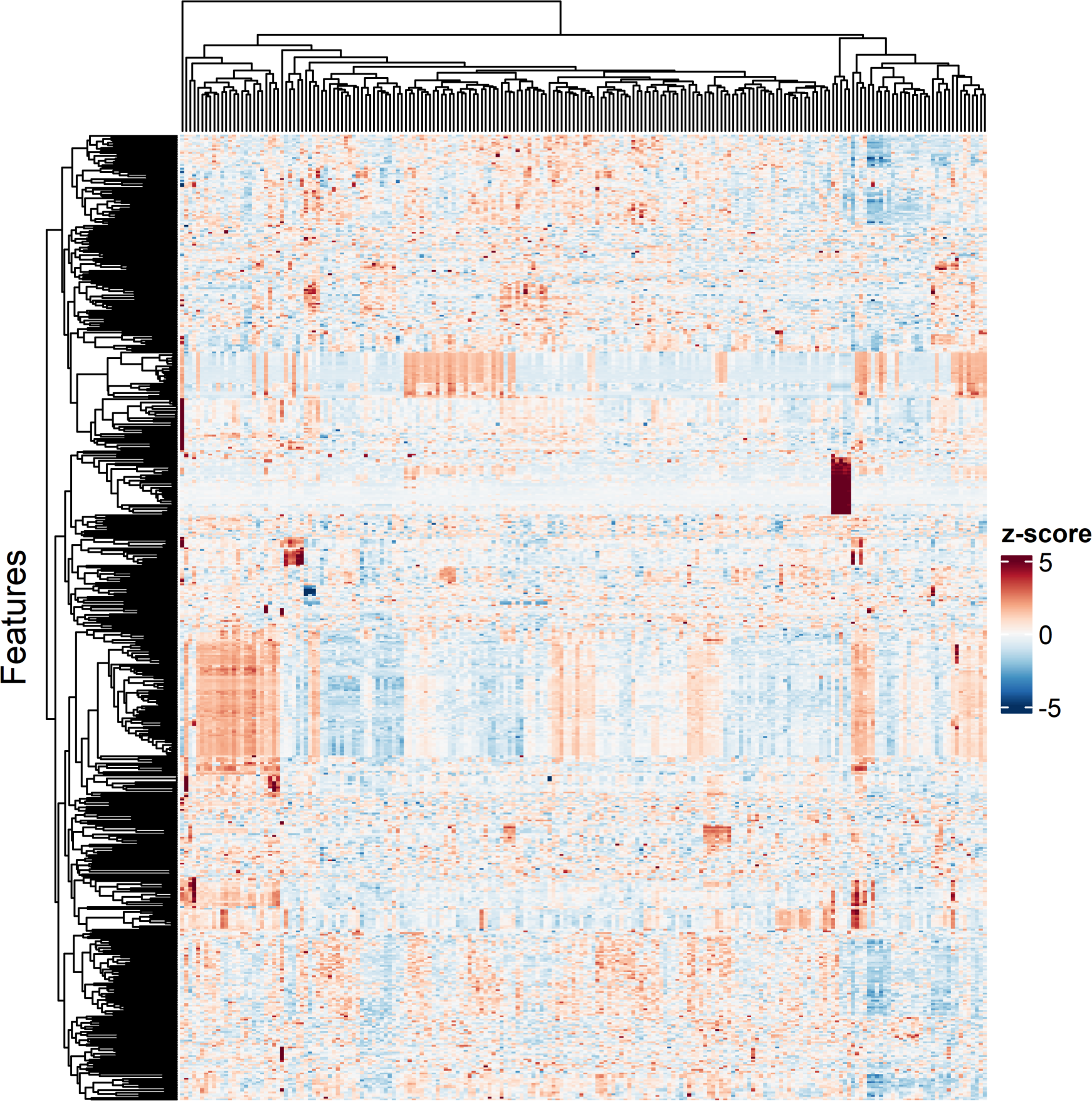
Heatmap representing untargeted metabolomic data of 202 leaf-derived protoplasts from the Sunstorm Apricot cultivar; 557 features were assigned a chemical formula (Supplementary Table 5, Supplementary Data 3).

**Supplementary Figure 7.**
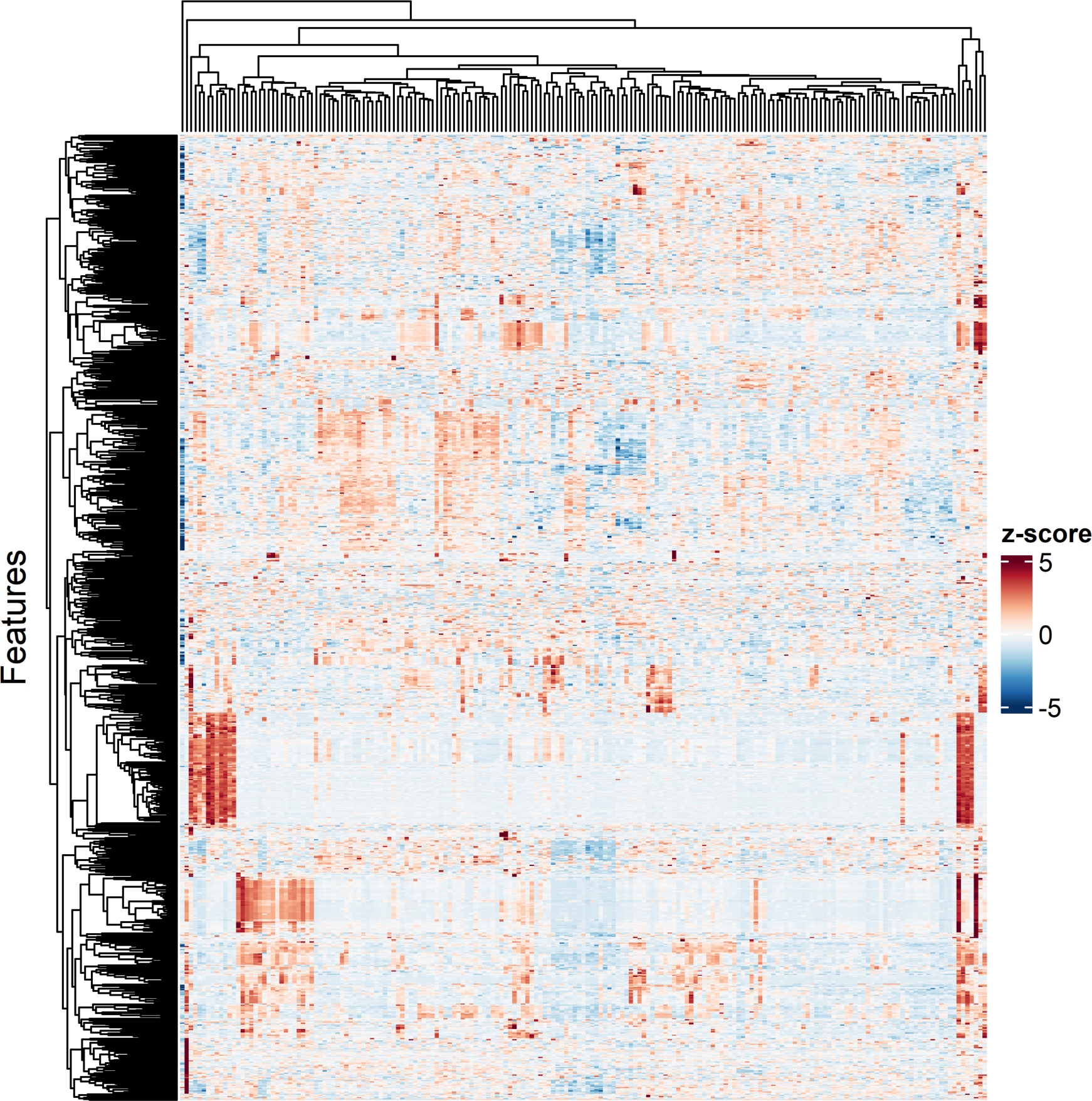
Heatmap representing untargeted metabolomic data of 187 root-derived protoplasts from the Sunstorm Apricot cultivar; 869 features were assigned a chemical formula (Supplementary Table 5, Supplementary Data 3).

**Supplementary Figure 8.**
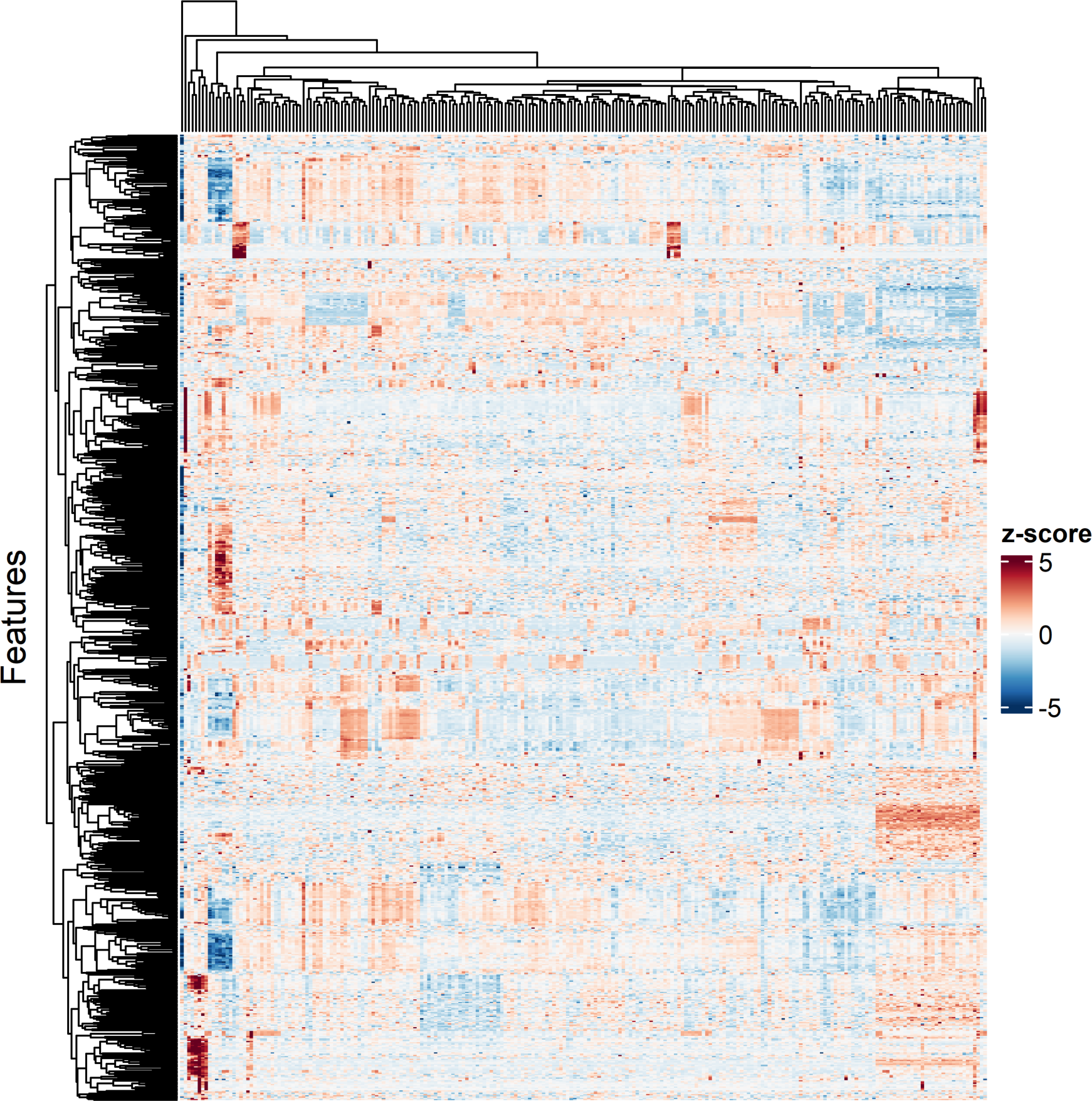
Heatmap representing untargeted metabolomic data of 232 petal-derived protoplasts from the Sunstorm Apricot cultivar; 765 features were assigned a chemical formula (Supplementary Table 5, Supplementary Data 3).

**Supplementary Figure 9.**
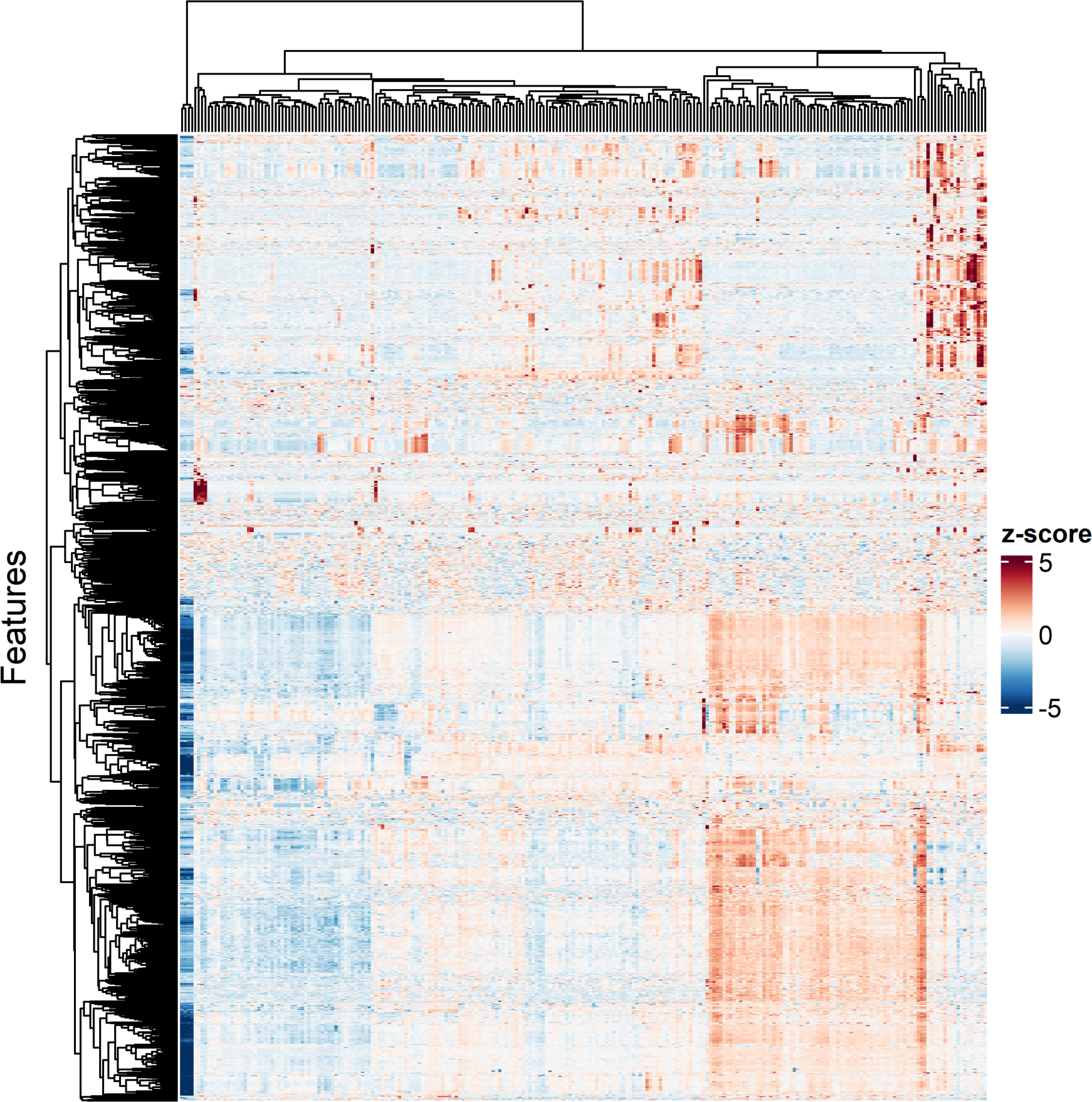
Heatmap representing untargeted metabolomic data of 241 petal-derived protoplasts from the Little Bright Eyes cultivar; 1373 features were assigned a chemical formula (Supplementary Table 5, Supplementary Data 3).

**Supplementary Figure 10.**
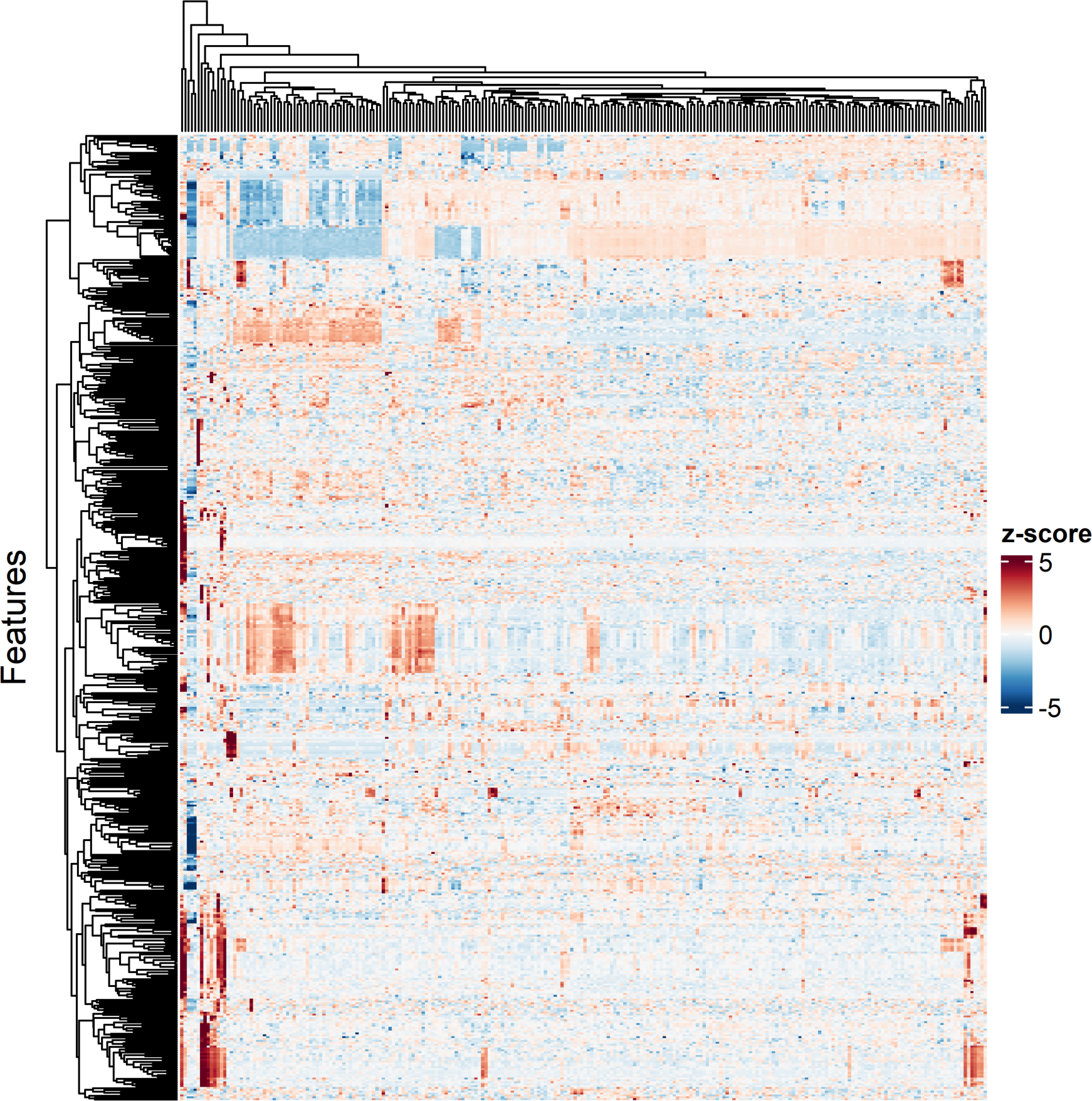
Heatmap representing untargeted metabolomic data of 244 petal-derived protoplasts from the Atlantic Burgundy Halo cultivar; 513 features were assigned a chemical formula (Supplementary Table 5, Supplementary Data 3).

**Supplementary Figure 11.**
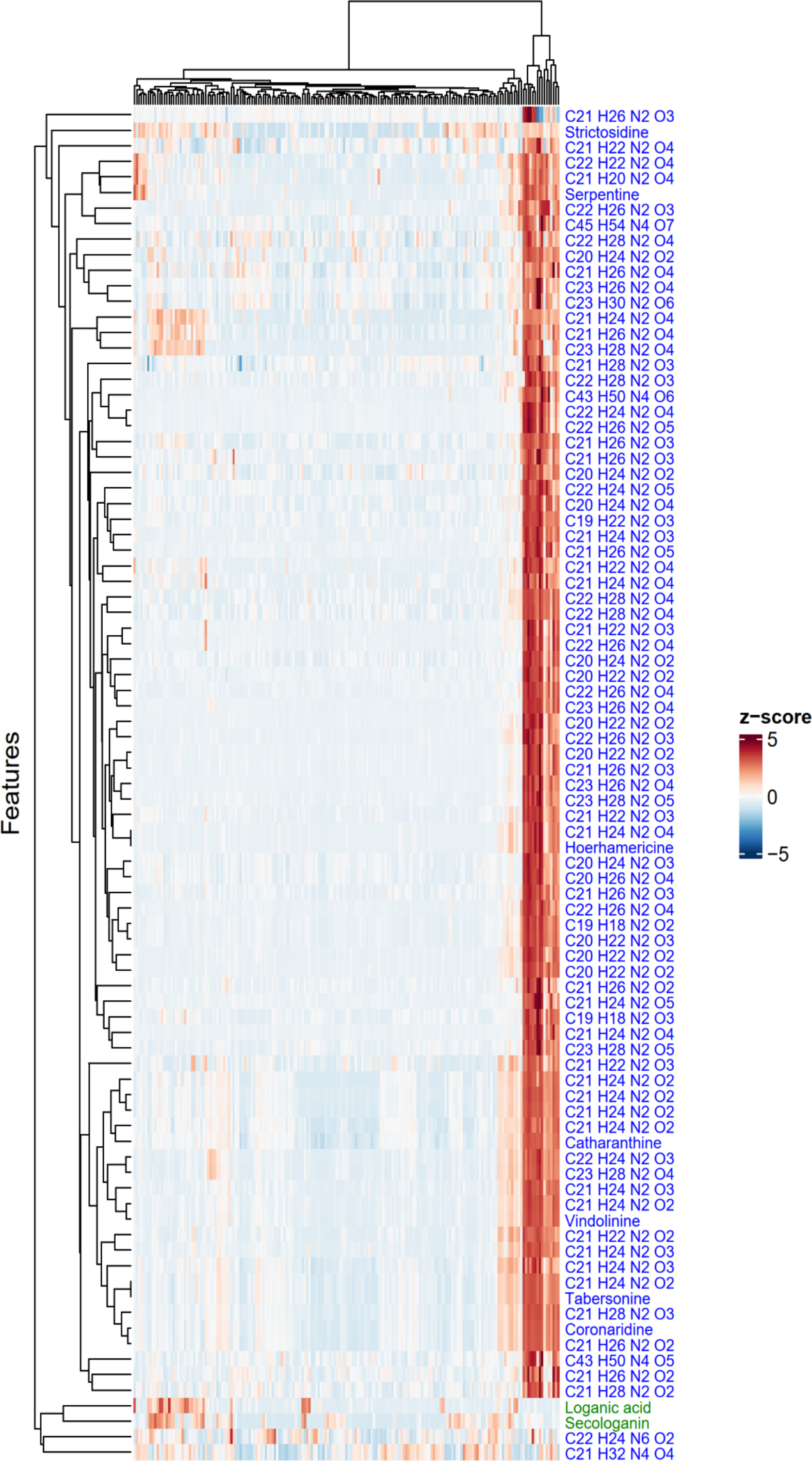
Heatmap showing the distribution of chemical features among the clusters. After analyzing 187 Sunstorm Apricot root protoplasts, 87 chemical features are confidently assigned to iridoid (2, in green), alkaloid (85, in blue) classes. Compounds for which a name is provided were structurally validated with an authentic standard (Supplementary Fig. 3) (See main text Fig. 3 for comparable data for SA leaf-derived protoplasts).

**Supplementary Figure 12.**
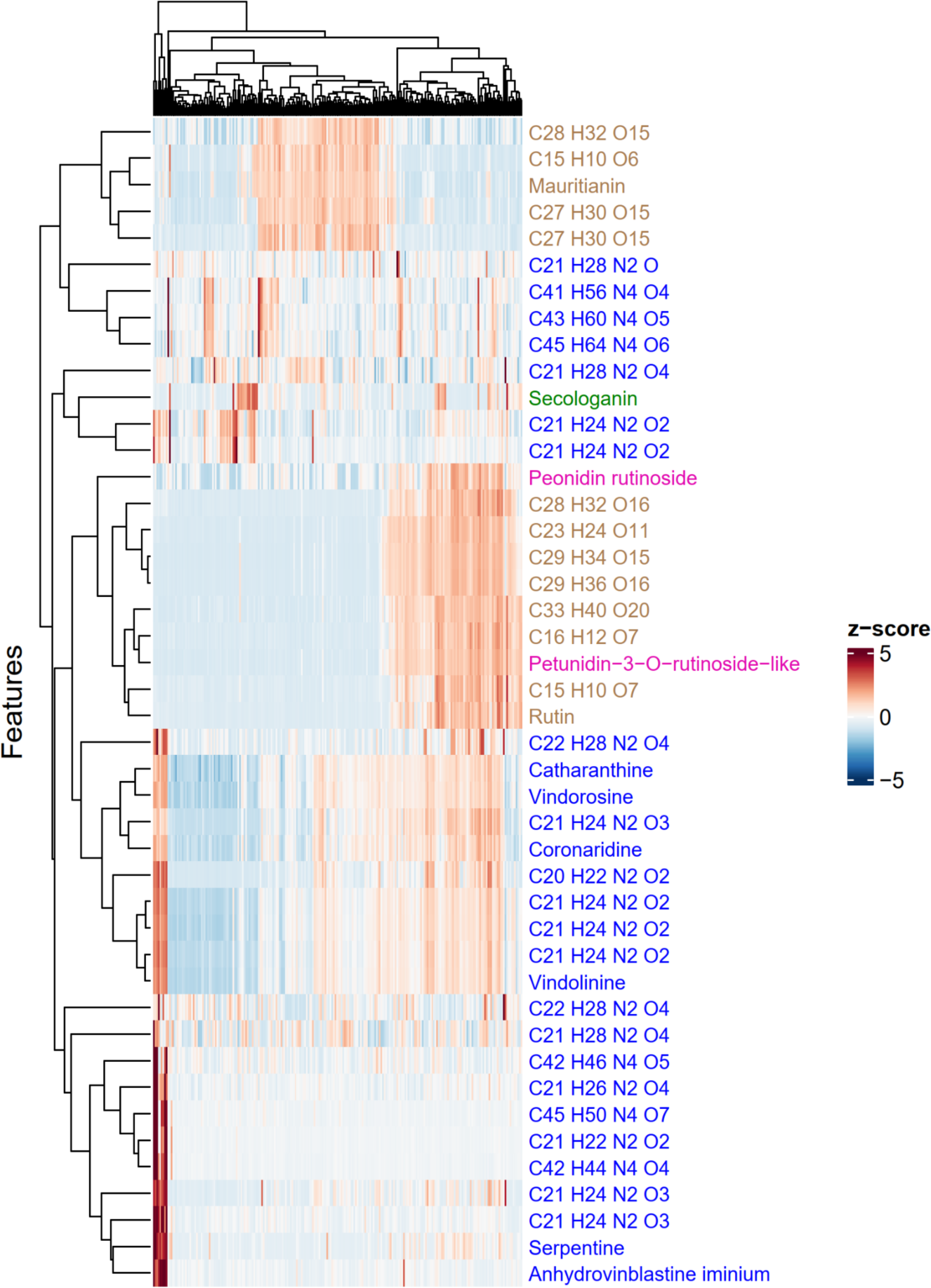
Heatmap showing the distribution of chemical features among the clusters. After analyzing 232 Sunstorm Apricot petal protoplasts, 44 chemical features are confidently assigned to iridoid (1, in green), alkaloid (28, in blue), flavonoid (13, in brown), or anthocyanin (2, in pink) classes. Compounds for which a name is provided were structurally validated with an authentic standard (Supplementary Fig. 3) (See main text Fig. 3 for comparable data for SA leaf-derived protoplasts).

**Supplementary Figure 13.**
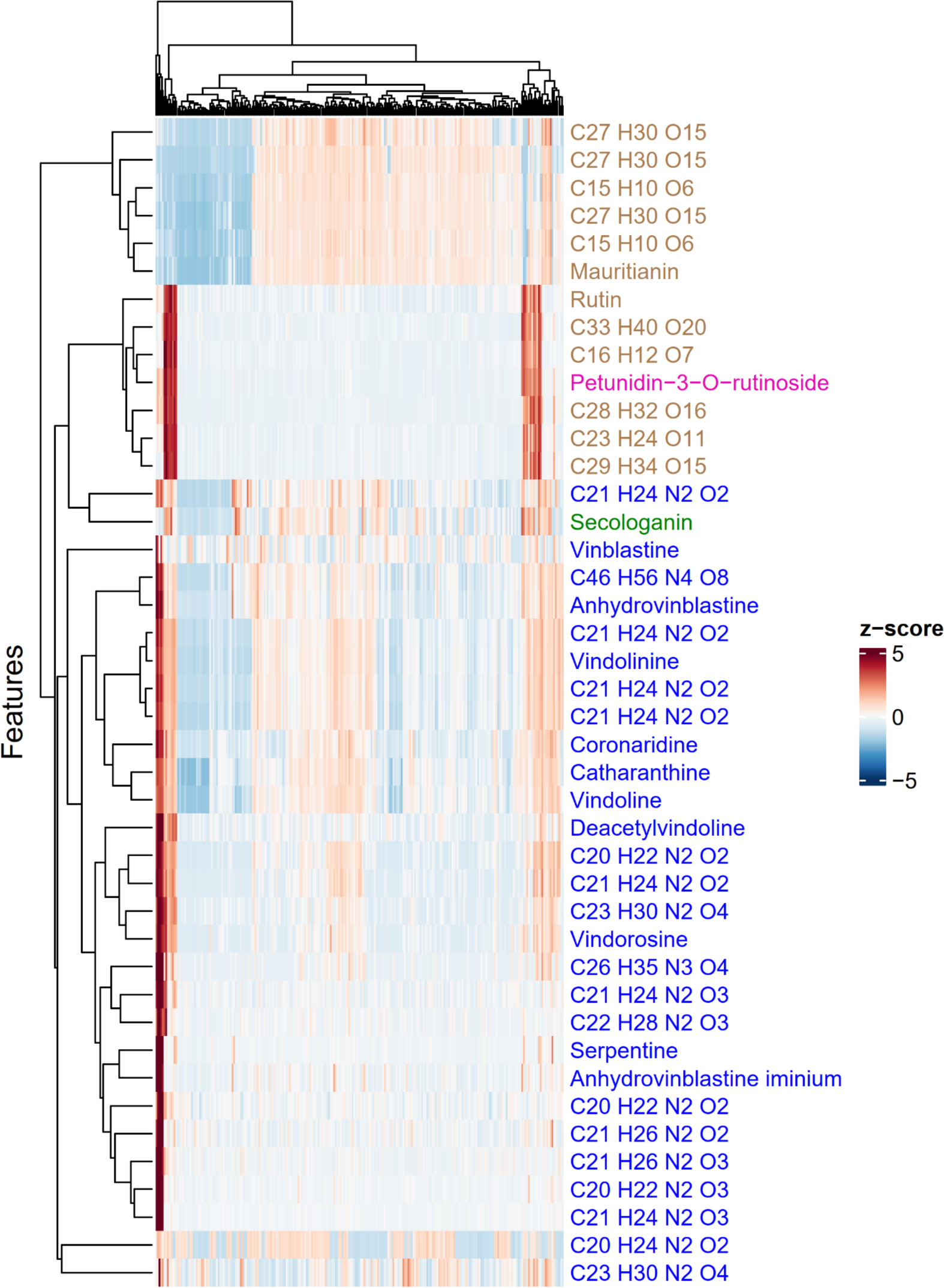
Heatmap showing the distribution of chemical features among the clusters. After analyzing 241 Little Bright Eyes petal protoplasts, 42 chemical features are confidently assigned to iridoid (1, in green), alkaloid (28, in blue), flavonoid (12, in brown), or anthocyanin (1, in pink) classes. Compounds for which a name is provided were structurally validated with an authentic standard (Supplementary Fig. 3) (See main text Fig. 3 for comparable data for SA leaf-derived protoplasts).

**Supplementary Figure 14.**
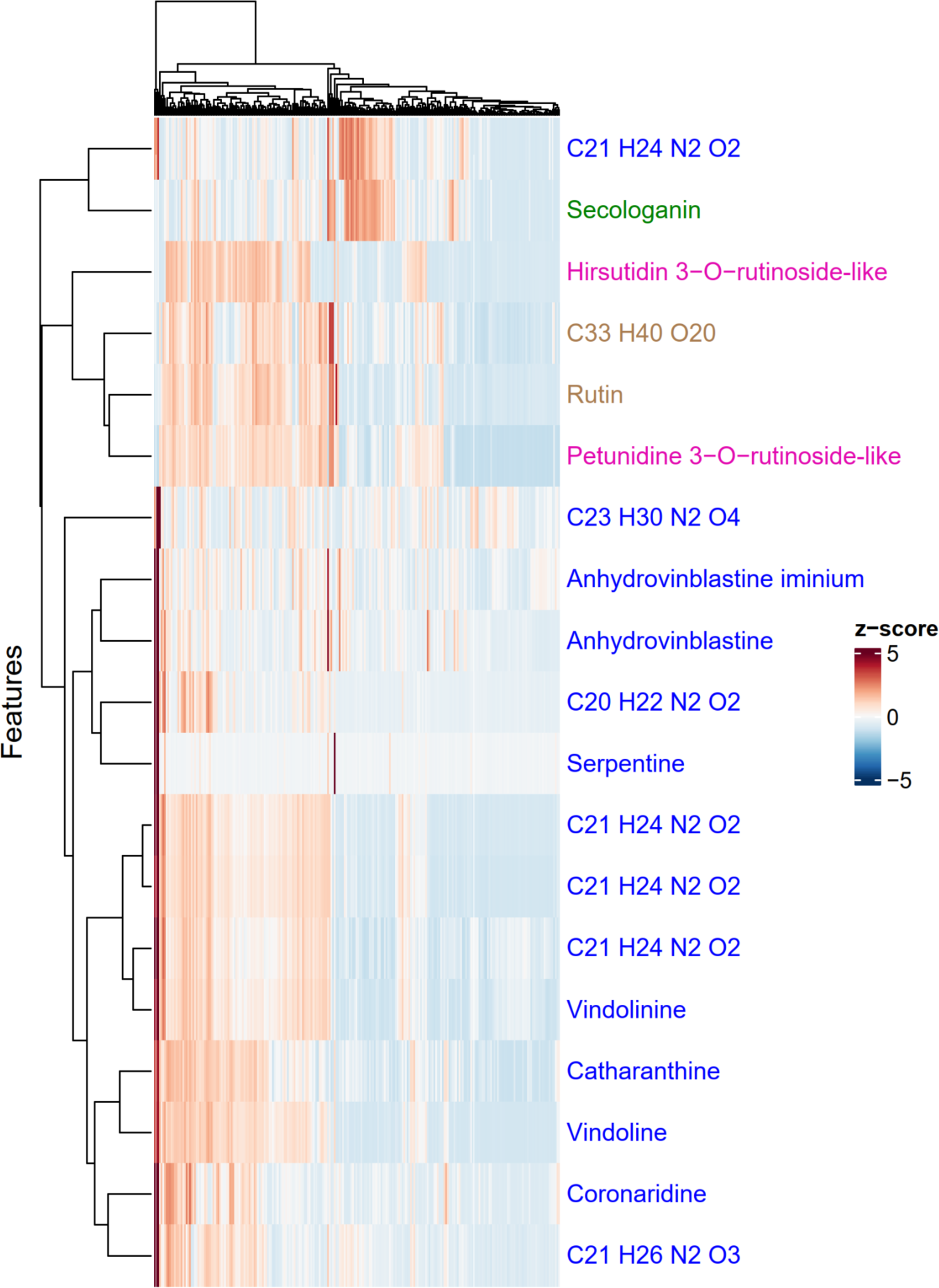
Heatmap showing the distribution of chemical features among the clusters. After analyzing 244 Atlantic Burgundy Halo petal protoplasts, 19 chemical features are confidently assigned to iridoid (1, in green), alkaloid (14, in blue), flavonoid (2, in brown), or anthocyanin (2, in pink) classes. Compounds for which a name is provided were structurally validated with an authentic standard (Supplementary Fig. 3) (See main text Fig. 3 for comparable data for SA leaf-derived protoplasts).

**Supplementary Figure 15.**
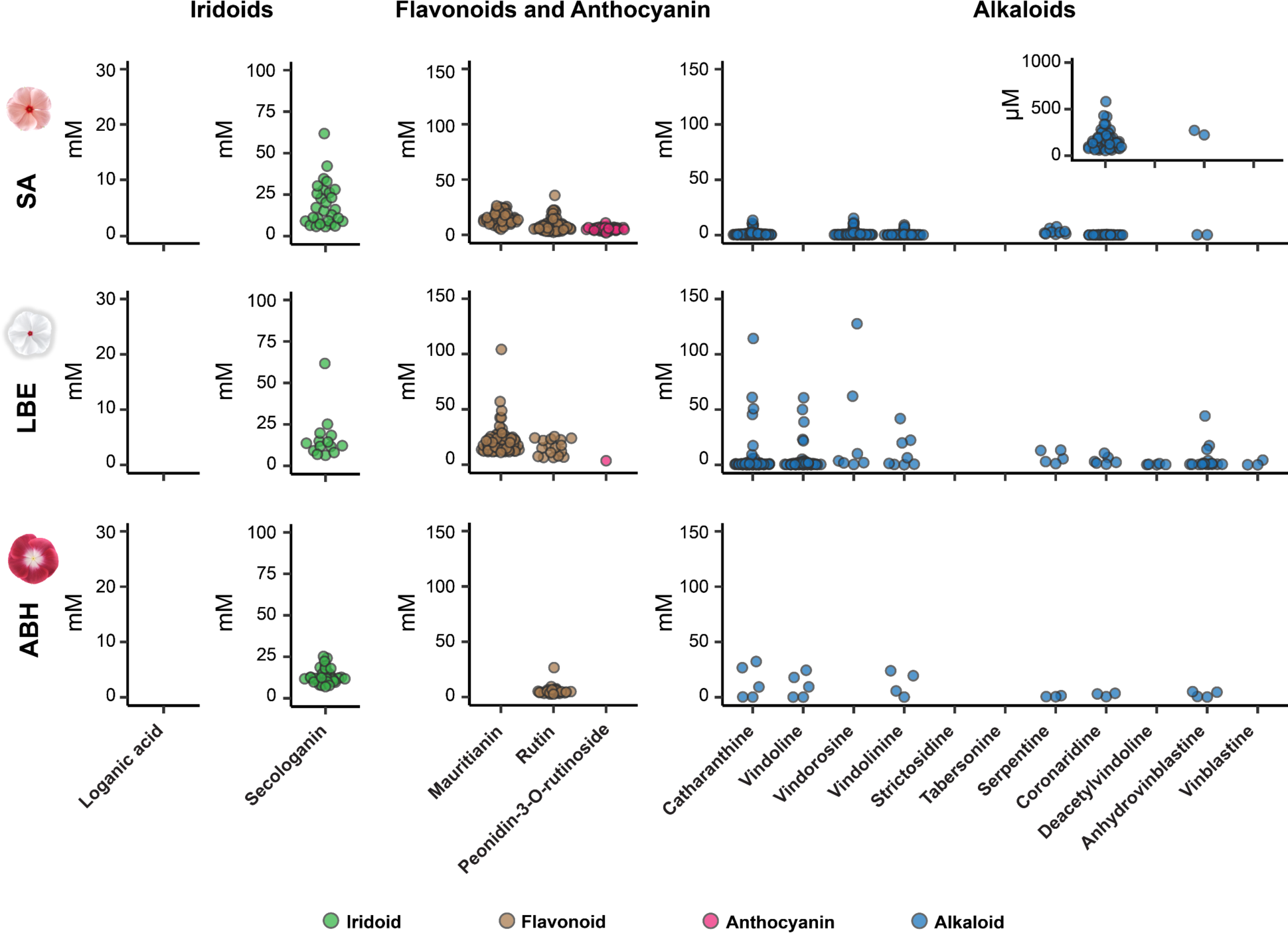
Quantification of iridoids, flavonoids, anthocyanins and alkaloids in single cells isolated from flower petals from three different varieties (Supplementary Data 4).

**Supplementary Figure 16.**
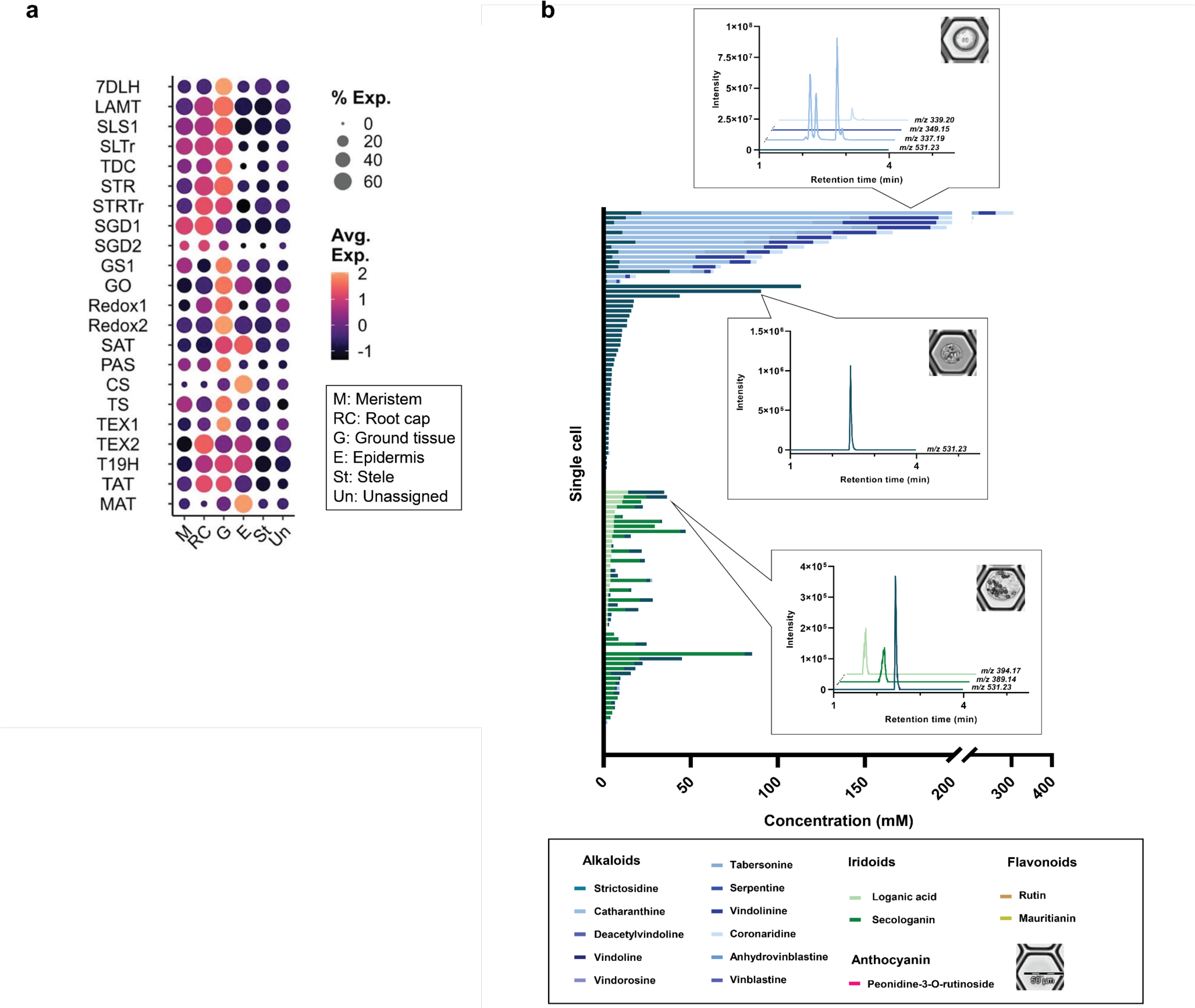
**a,** Single cell mRNA data of root protoplasts (Sunstorm Apricot cultivar). Data are taken from Li et al. 2023^1^. Dotplot shows cell-type specific expression patterns of MIA biosynthesis (See Supplementary Table 1 for definition of enzyme abbreviations). **b,** Ratio of compounds found across the population of root cells that were measured (187 cells). Stack plot showing the absolute concentration of each of the quantified metabolites in each cell. Colors indicate classes of compounds: blue bars represent alkaloids, dark green represents secologain (iridoid), light green represents loganic acid (iridoid). Representative chromatograms of these compounds from individual cells. *m/z* 394.17, loganic acid; *m/z* 389.14, secologanin; *m/z* 531.23, strictosidine; *m/z* 337.19, catharanthine and vindolinine; *m/z* 339.20, coronaridine; *m/z* 349.15, serpentine (See main text Fig. 5 for comparable data for SA leaf-derived protoplasts) (Supplementary Data 4).

**Supplementary Figure 17.**
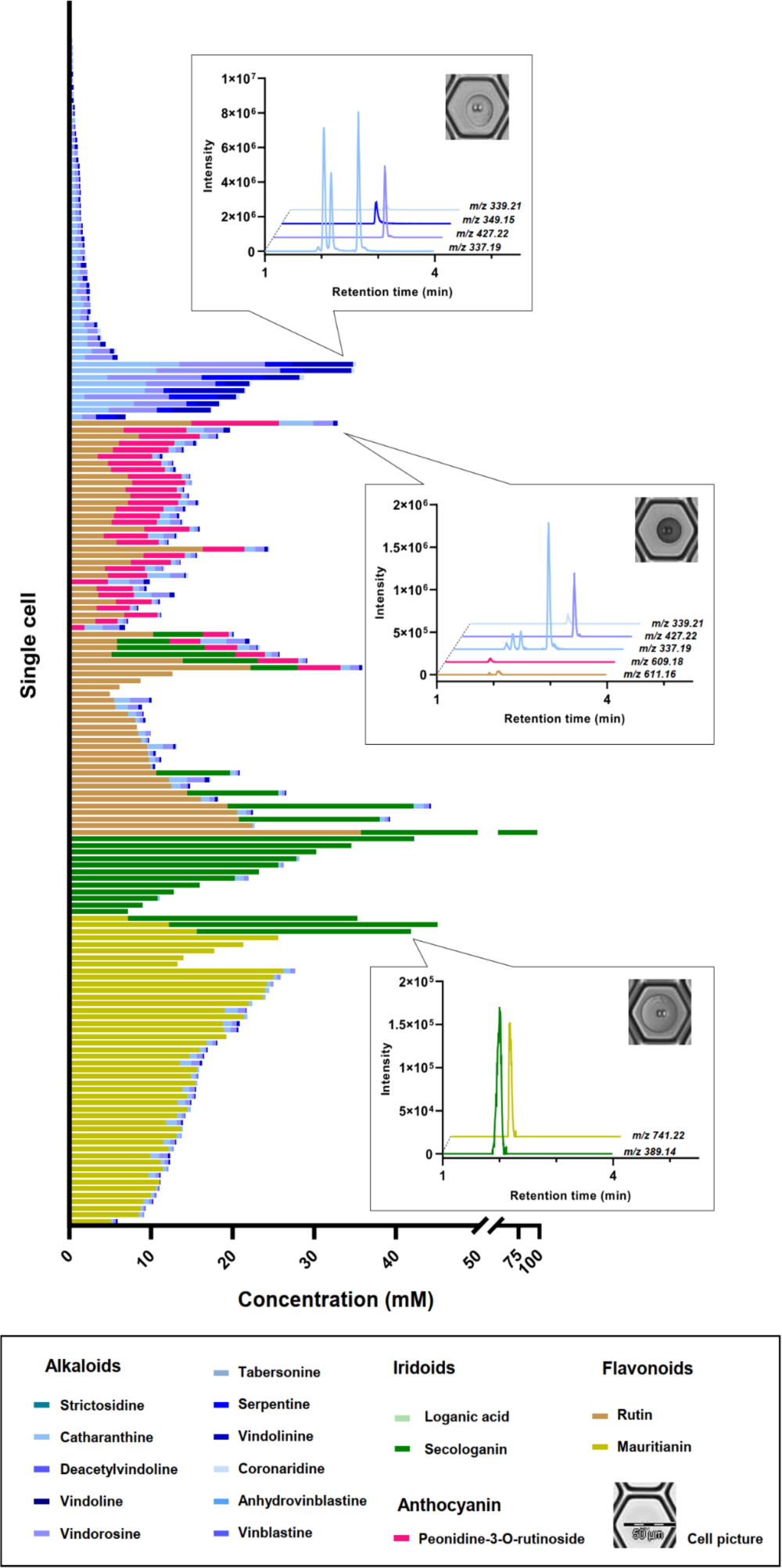
Ratio of compounds found across the population of petal cells (Sunstorm Apricot cultivar) that were measured (232 cells). Stack plot showing the absolute concentration of each of the quantified metabolites in each cell. Colors indicate classes of compounds: blue bars represent alkaloids, dark green represents secologain (iridoid), light green represents loganic acid (iridoid), mustard represents mauritanin (flavonoid), brown represents rutin (flavonoid) and pink represents peonidin 3-O-rutinoside (anthocyanin). Representative chromatograms of these compounds from individual cells. *m/z* 389.14, secologanin; *m/z* 741.22, mauritanin; *m/z* 611.16, rutin; *m/z* 609.18, peonidin 3-O-rutinoside; *m/z* 337.19, catharanthine and vindolinine; *m/z* 339.21, coronaridine; *m/z* 427.22, vindorosine; *m/z* 349.15, serpentine (See main text Fig. 5 for comparable data for SA leaf-derived protoplasts) (Supplementary Data 4).

**Supplementary Figure 18.**
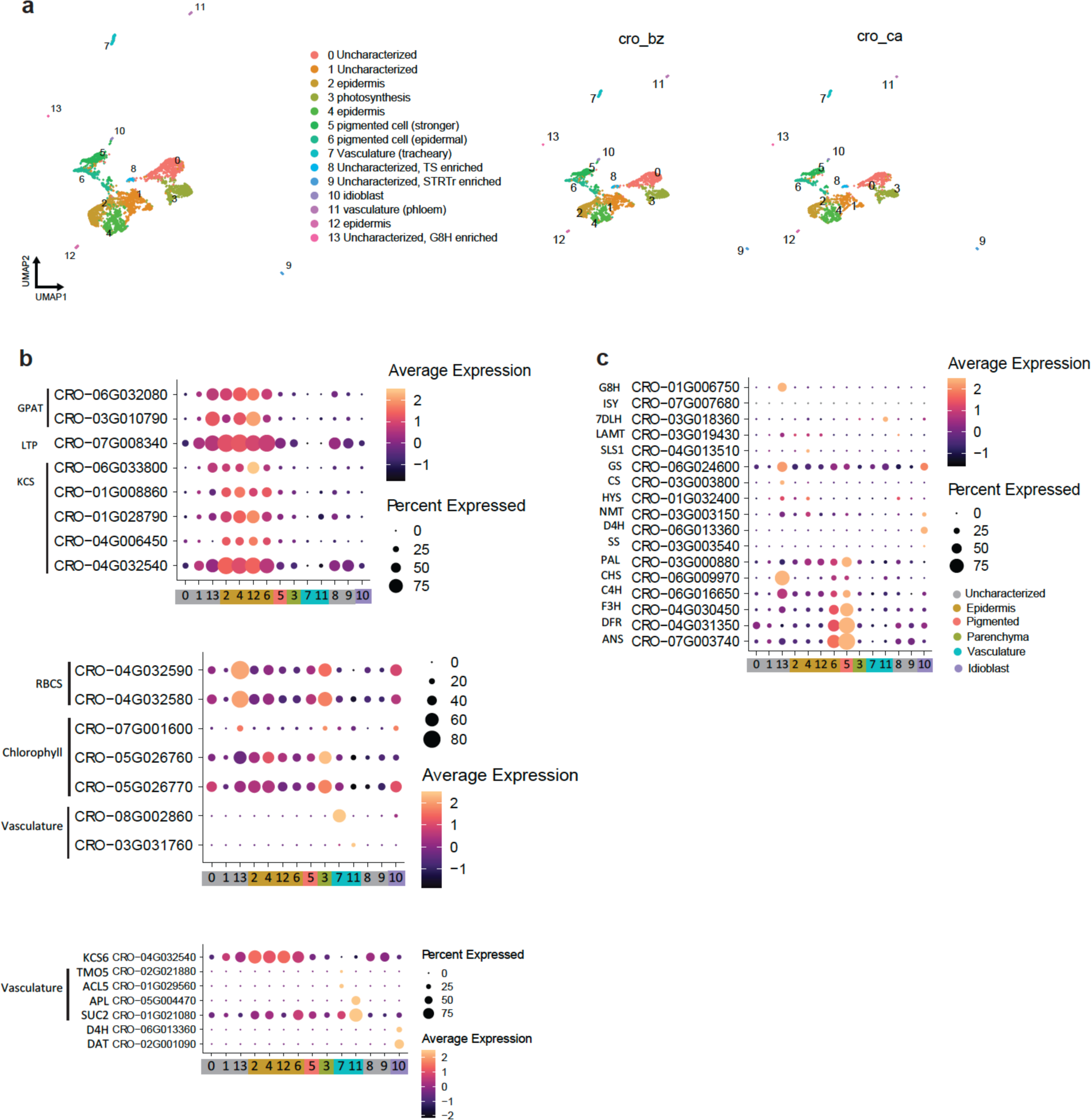
Single cell mRNA data of Sunstorm Apricot (SA) petals. **a**, UMAP plot of scRNA-Seq for SA petals. **b**, Dotplots for marker genes. Cuticle biosynthesis genes (GPAT, LTP, KCS), chloroplast-related genes (RBCS, Chlorophyll binding proteins), vasculature-related genes were identified as de novo marker genes for the clusters. Marker genes previously used in leaf datasets (Li et al. 2023^1^) also confirmed the cell annotations. **c,** Dotplot shows cell-type specific expression patterns of MIA biosynthetic enzymes (See Supplementary Table 1 for definition of enzyme abbreviations) (Supplementary Data 5).

**Supplementary Figure 19.**
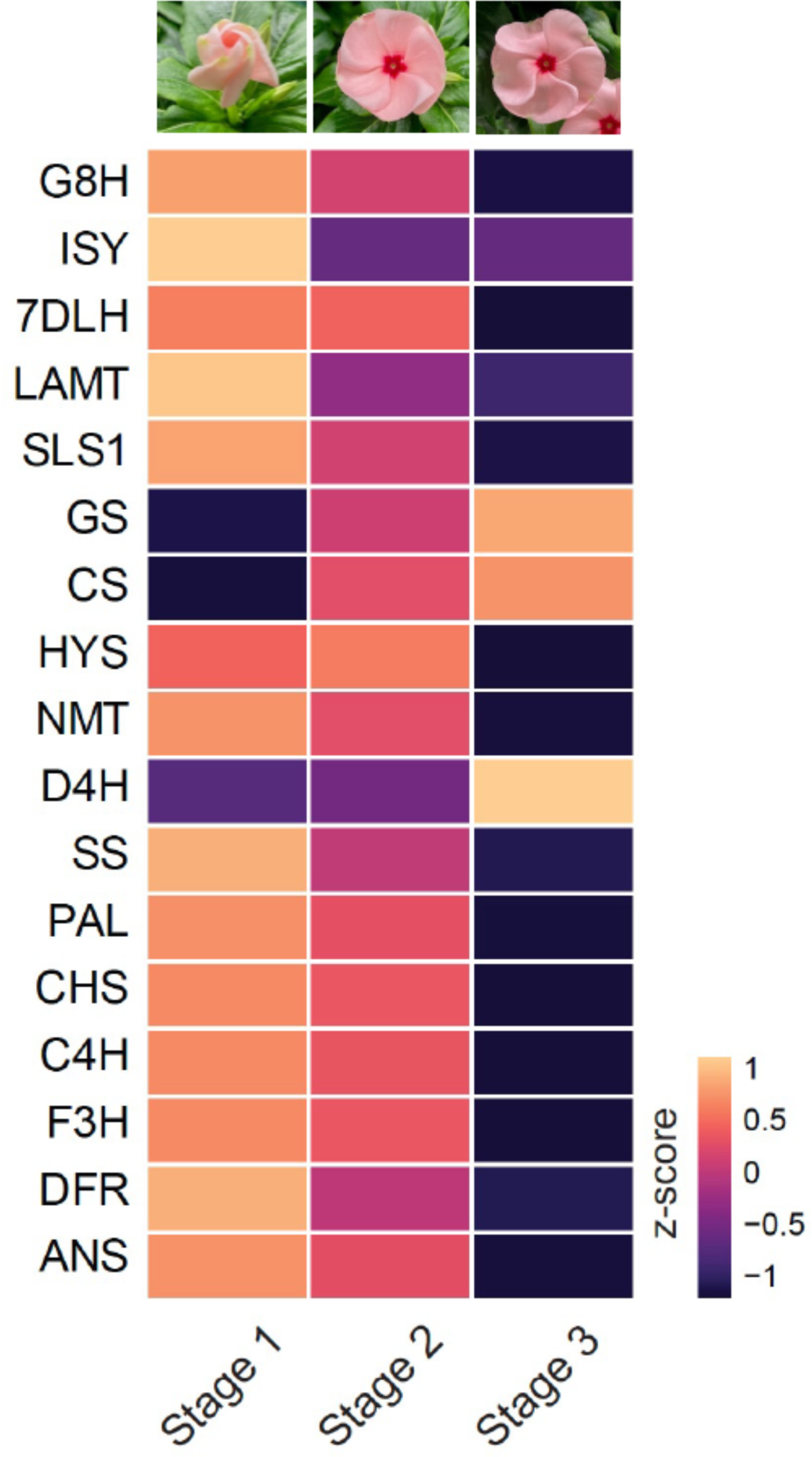
Bulk RNAseq data at a variety of time points immediately after Sunstorm Apricot flower opening (See Supplementary Table 1 for definition of enzyme abbreviations) (Supplementary Data 6).

**Supplementary Figure 20.**
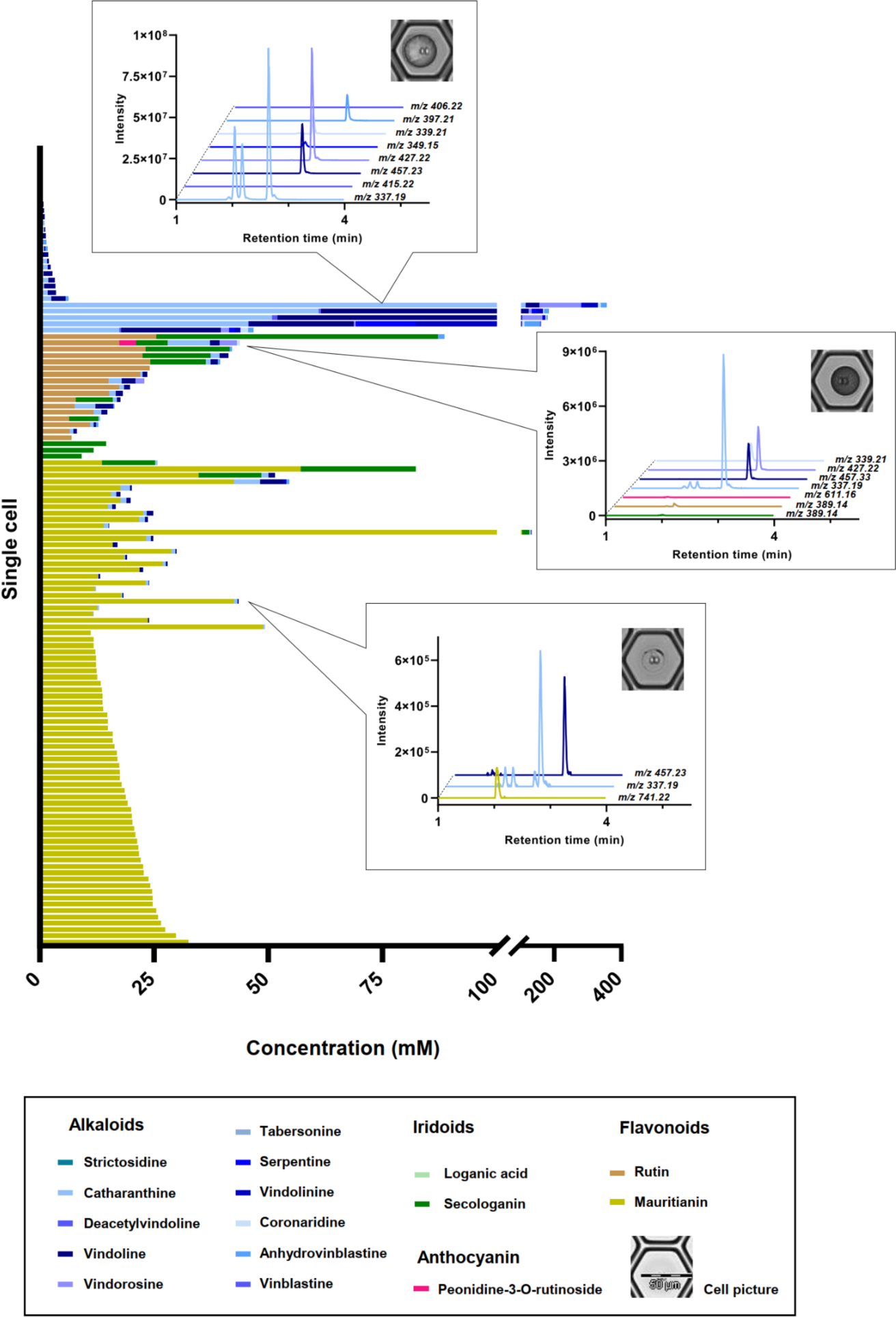
Ratio of compounds found across the population of petal cells (Little Bright Eyes cultivar) that were measured (241 cells). Stack plot showing the absolute concentration of each of the quantified metabolites in each cell. Colors indicate classes of compounds: blue bars represent alkaloids, dark green represents secologain (iridoid), mustard represents mauritanin (flavonoid), brown represents rutin (flavonoid) and pink represents peonidin 3-O-rutinoside (anthocyanin). Representative chromatograms of these compounds from individual cells. *m/z* 389.14, secologanin; *m/z* 389.14, rutin; *m/z* 741.22, mauritanin; *m/z* 611.16, peonidin-3-O-rutinoside; *m/z* 337.19, catharanthine and vindolinine; *m/z* 457.23, vindoline; *m/z* 427.22, vindorosine; *m/z* 349.15, serpentine; *m/z* 415.22, deacetylvindoline; *m/z* 339.21, coronaridine; *m/z* 397.21, anhydrovinblastine; *m/z* 406.22, vinblastine (See main text Fig. 5 for comparable data for SA leaf-derived protoplasts) (Supplementary Data 4).

**Supplementary Figure 21.**
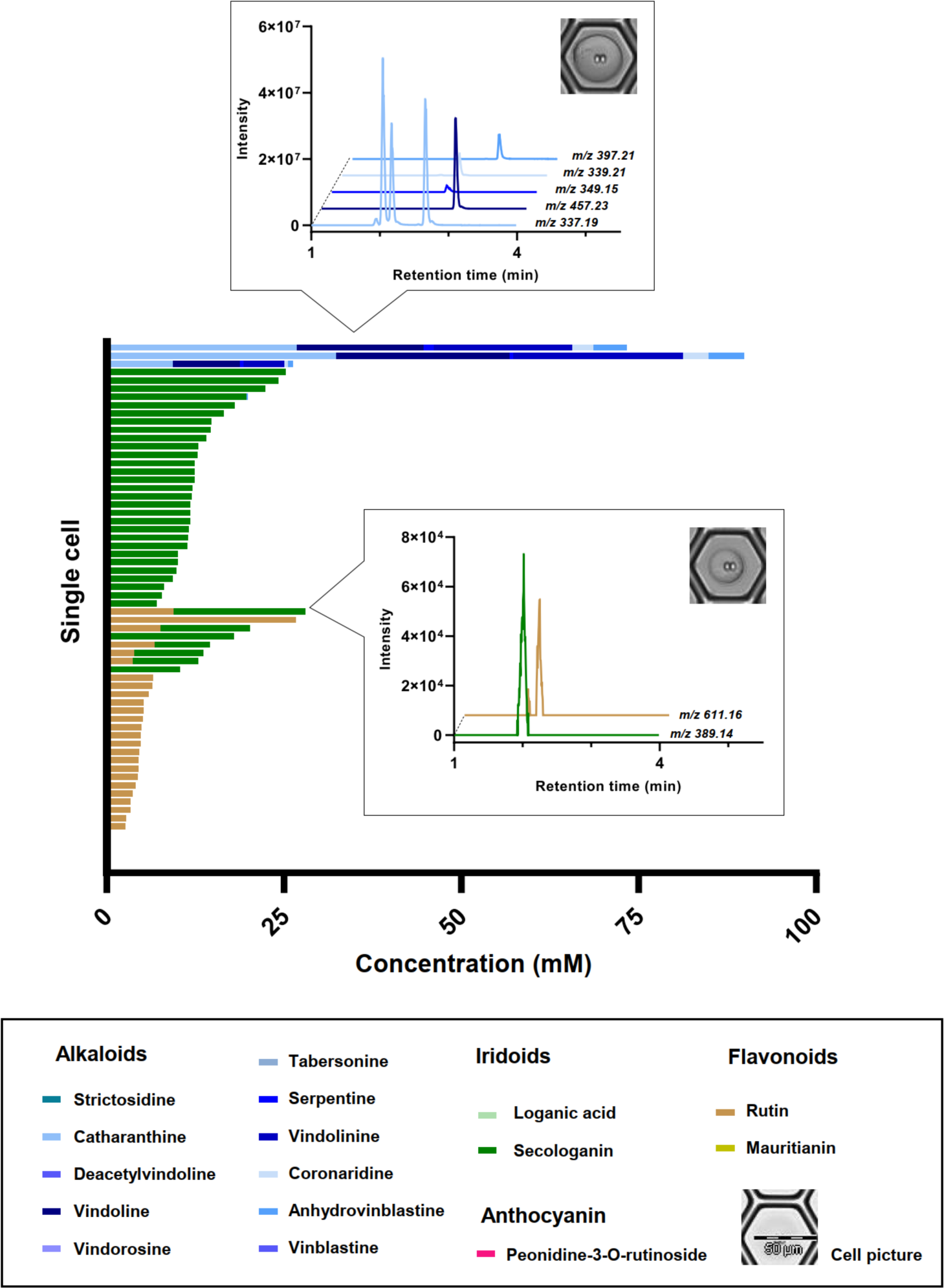
Ratio of compounds found across the population of petal cells (Atlantic Burgundy Halo cultivar) that were measured (244 cells). Stack plot showing the absolute concentration of each of the quantified metabolites in each cell. Colors indicate classes of compounds: blue bars represent alkaloids, dark green represents secologain (iridoid), light green represents loganic acid (iridoid), mustard represents mauritanin (flavonoid), brown represents rutin (flavonoid) and pink represents penodine-3-O-rutinoside (anthocyanin). Representative chromatograms of these compounds from individual cells. *m/z* 389.14, secologanin; *m/z* 611.16, rutin; *m/z* 337.19, catharanthine and vindolinine; *m/z* 457.23, vindoline; *m/z* 349.15, serpentine; *m/z* 397.21, anhydrovinblastine (See main text Fig. 5 for comparable data for SA leaf-derived protoplasts) (Supplementary Data 4).

**Supplementary Figure 22.**
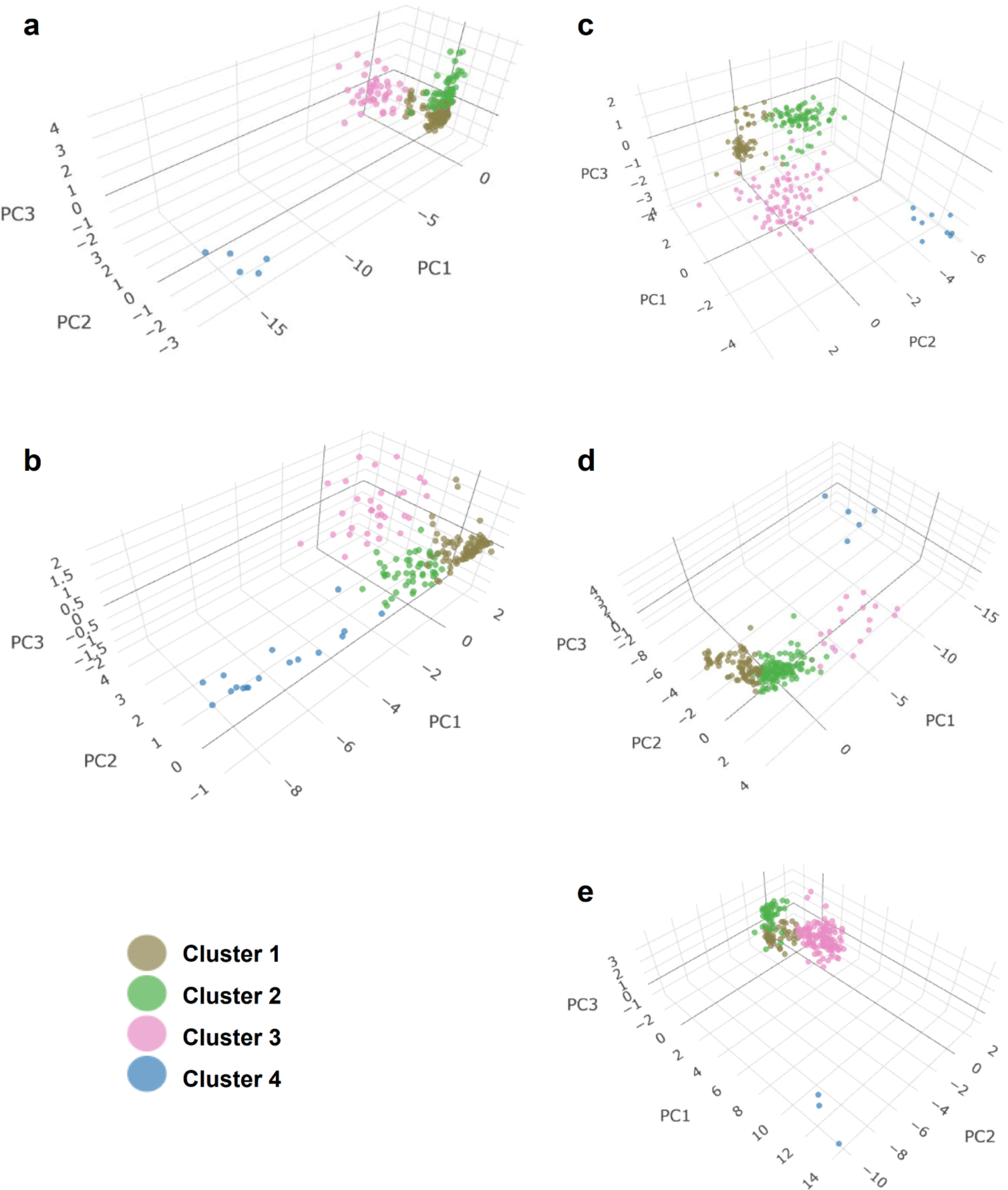
PCA analysis of single cells using only the compounds identified by co-elution with a reference standard from 5 different tissues: a. Sunstorm Apricot leaf, **b,** Sunstorm Apricot root, **c**, Sunstorm Apricot petal, **d**, Little Bright Eyes petal, **e,** Atlantic Burgundy Halo petal. Using k-means clustering analysis, the optimal number of clusters was defined as 4 using the Elbow method^2^. Using these parameters, cells were classified into four well-defined and distinct clusters.

### Supplementary Tables

**Supplementary Table 1.**
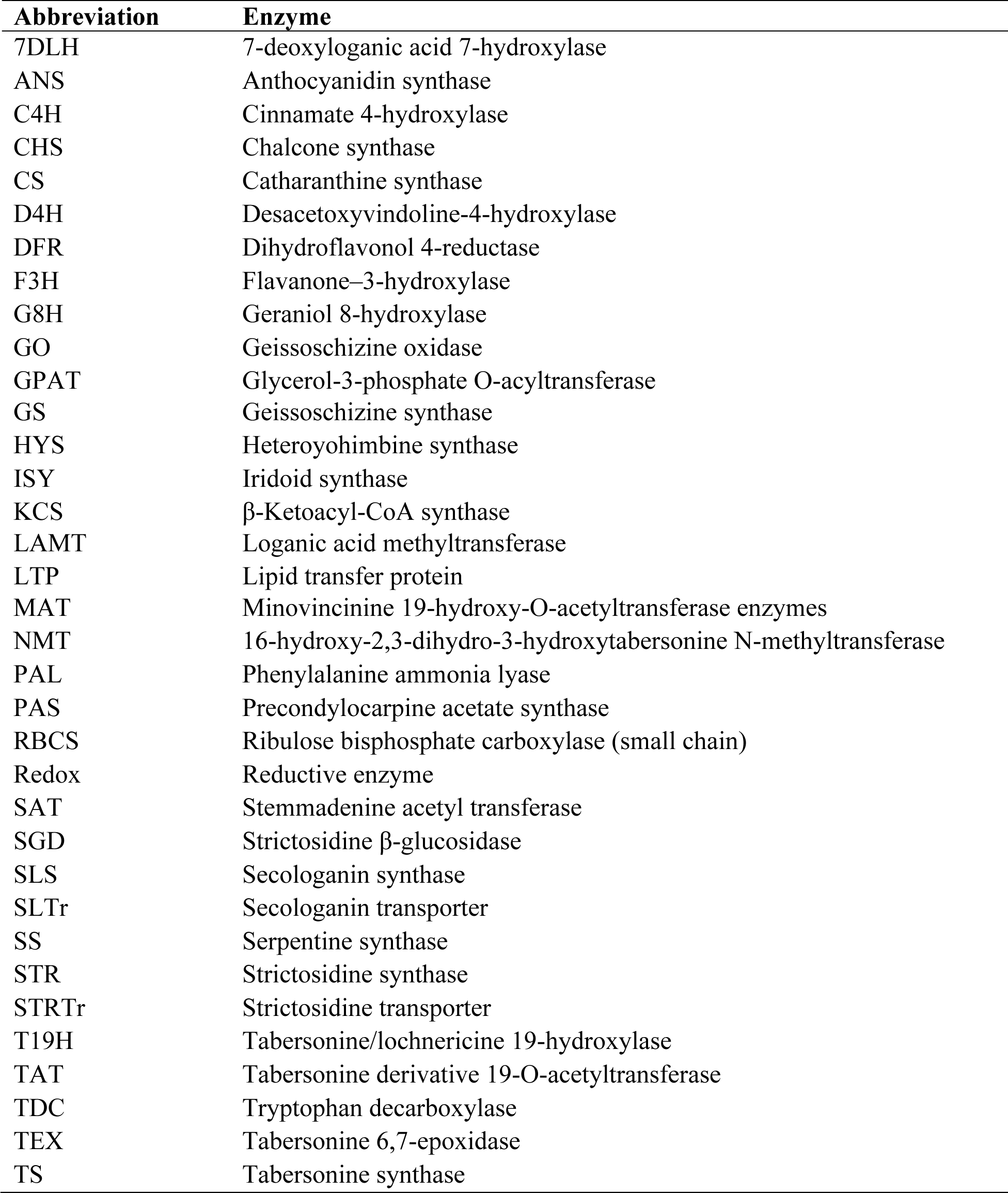
Definition of enzyme abbreviations.

| Abbreviation | Enzyme |
| --- | --- |
| 7DLH | 7-deoxyloganic acid 7-hydroxylase |
| ANS | Anthocyanidin synthase |
| C4H | Cinnamate 4-hydroxylase |
| CHS | Chalcone synthase |
| CS | Catharanthine synthase |
| D4H | Desacetoxyvindoline-4-hydroxylase |
| DFR | Dihydroflavonol 4-reductase |
| F3H | Flavanone-3-hydroxylase |
| G8H | Geraniol 8-hydroxylase |
| GO | Geissoschizine oxidase |
| GPAT | Glycerol-3-phosphate O-acyltransferase |
| GS | Geissoschizine synthase |
| HYS | Heteroyohimbine synthase |
| ISY | Iridoid synthase |
| KCS | β-Ketoacyl-CoA synthase |
| LAMT | Loganic acid methyltransferase |
| LTP | Lipid transfer protein |
| MAT | Minovincinine 19-hydroxy-O-acetyltransferase enzymes |
| NMT | 16-hydroxy-2,3-dihydro-3-hydroxytabersonine N-methyltransferase |
| PAL | Phenylalanine ammonia lyase |
| PAS | Precondylocarpine acetate synthase |
| RBCS | Ribulose biphosphate carboxylase (small chain) |
| Redox | Reductive enzyme |
| SAT | Stemmadenine acetyl transferase |
| SGD | Strictosidine β-glucosidase |
| SLS | Secologanin synthase |
| SLTr | Secologanin transporter |
| SS | Serpentine synthase |
| STR | Strictosidine synthase |
| STRTr | Strictosidine transporter |
| T19H | Tabersonine/lochnericine 19-hydroxylase |
| TAT | Tabersonine derivative 19-O-acetyltransferase |
| TDC | Tryptophan decarboxylase |
| TEX | Tabersonine 6,7-epoxidase |
| TS | Tabersonine synthase |

**Supplementary Table 2.**
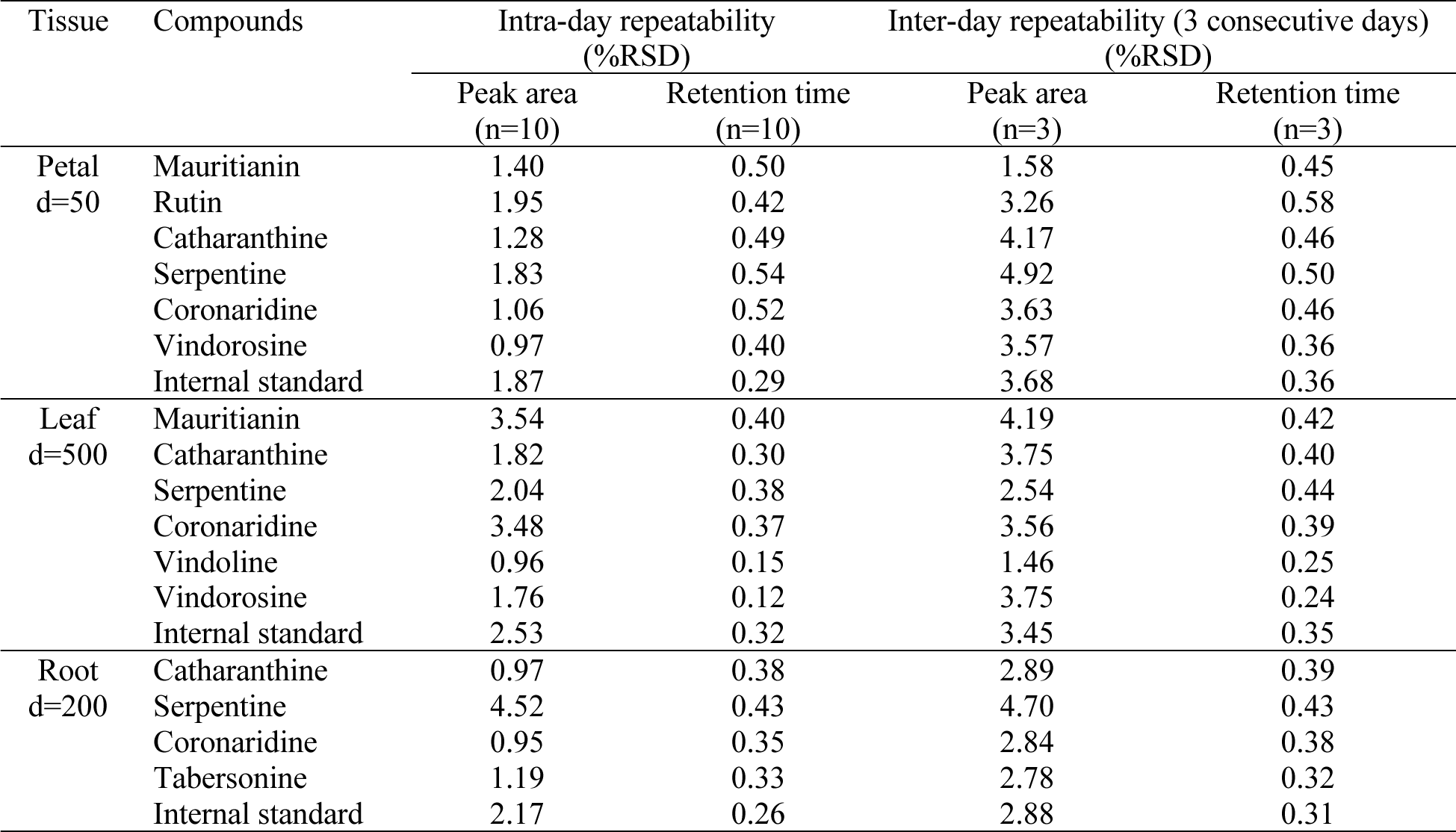
Validation of the reproducibility of the mass spectral data. Results are presented as RSD% of the peak area and retention time.

**Supplementary Table 3.** Compounds used for identification.

| Compound class | Compound name | Chemical formula | Ion | Measured m/z | Δppm | Retention time (min) |
| --- | --- | --- | --- | --- | --- | --- |
| Iridoids | Loganic acid | C<sub>16</sub>H<sub>24</sub>O<sub>10</sub> | [M+NH<sub>4</sub>]<sup>+</sup> | 394.17062 | 0.38 | 1.48 |
|  | Secologanin | C<sub>17</sub>H<sub>24</sub>O<sub>10</sub> | [M+H]<sup>+</sup> | 389.14429 | 0.18 | 2.02 |
| Flavonoids | Mauritianin | C<sub>33</sub>H<sub>40</sub>O<sub>19</sub> | [M+H]<sup>+</sup> | 741.22357 | 0.12 | 2.09 |
|  | Rutin | C<sub>27</sub>H<sub>30</sub>O<sub>16</sub> | [M+H]<sup>+</sup> | 611.16064 | 0.03 | 2.08 |
| Anthocyanins | Peonidin 3-O-rutinoside | C<sub>28</sub>H<sub>33</sub>O<sub>15</sub> | [M]<sup>+</sup> | 609.18164 | 0.39 | 1.80 |
|  | Petunidin 3-O-rutinoside* | C<sub>28</sub>H<sub>33</sub>O<sub>16</sub> | [M]<sup>+</sup> | 625.17554 | 1.23 | 2.30 |
|  | Hirsutidin 3-O-rutinoside* | C<sub>30</sub>H<sub>37</sub>O<sub>16</sub> | [M]<sup>+</sup> | 653.20673 | 1.35 | 2.04 |
| Alkaloids | Strictosidine** | C<sub>27</sub>H<sub>34</sub>N<sub>2</sub>O<sub>9</sub> | [M+H]<sup>+</sup> | 531.23383 | 0.24 | 2.43 |
|  | Catharanthine | C<sub>21</sub>H<sub>24</sub>N<sub>2</sub>O<sub>2</sub> | [M+H]<sup>+</sup> | 337.19064 | 1.22 | 2.65 |
|  | Deacetylvindoline | C<sub>23</sub>H<sub>30</sub>N<sub>2</sub>O<sub>5</sub> | [M+H]<sup>+</sup> | 415.22247 | 0.67 | 2.58 |
|  | Vindoline | C<sub>25</sub>H<sub>32</sub>N<sub>2</sub>O<sub>6</sub> | [M+H]<sup>+</sup> | 457.23325 | 0.13 | 2.97 |
|  | Tabersonine | C<sub>21</sub>H<sub>24</sub>N<sub>2</sub>O<sub>2</sub> | [M+H]<sup>+</sup> | 337.19080 | 0.74 | 2.77 |
|  | Hörhammericine** | C<sub>21</sub>H<sub>24</sub>N<sub>2</sub>O<sub>4</sub> | [M+H]<sup>+</sup> | 369.18051 | 0.79 | 1.77 |
|  | Serpentine | C<sub>21</sub>H<sub>20</sub>N<sub>2</sub>O<sub>3</sub> | [M]<sup>+</sup> | 349.15448 | 0.54 | 2.65 |
|  | Vindolinine | C<sub>21</sub>H<sub>24</sub>N<sub>2</sub>O<sub>2</sub> | [M+H]<sup>+</sup> | 337.19077 | 0.83 | 2.19 |
|  | Coronaridine** | C<sub>21</sub>H<sub>26</sub>N<sub>2</sub>O<sub>2</sub> | [M+H]<sup>+</sup> | 339.20657 | 0.38 | 2.70 |
|  | Vindorosine** | C<sub>24</sub>H<sub>30</sub>N<sub>2</sub>O<sub>5</sub> | [M+H]<sup>+</sup> | 427.22241 | 0.80 | 2.99 |
|  | Anhydrovinblastine iminium** | C<sub>46</sub>H<sub>54</sub>N<sub>4</sub>O<sub>8</sub> | [M+2H]<sup>2+</sup> | 396.20432 | 0.10 | 3.00 |
|  | Anhydrovinblastine | C<sub>46</sub>H<sub>56</sub>N<sub>4</sub>O<sub>8</sub> | [M+2H]<sup>2+</sup> | 397.21222 | 0.10 | 3.16 |
|  | Vinblastine | C<sub>46</sub>H<sub>58</sub>N<sub>4</sub>O<sub>9</sub> | [M+2H]<sup>2+</sup> | 406.21741 | 0.15 | 2.94 |

\*Putative assignment based on mass and fragmentation of other glucosides with the same aglycon, standard is not available.
\*\* Standards were synthesized/isolated in our laboratory.

**Supplementary Table 4.** Analytical parameters of the compounds quantified in this study.

| Compound | Calibration range (nM) | Regression equation<sup>a</sup> | Correlation Coefficient (R<sup>2</sup>) | LOQ<sup>b</sup> (nM) |
| --- | --- | --- | --- | --- |
| Loganic acid | 3.5-9000 | y=16211x + 242751 | 0.9999 | 3.5 |
| Secologanin | 8.0-3000 | y = 13483x + 429913 | 0.9987 | 8 |
| Mauritianin | 10.0-3000 | y = 13958x + 42965 | 0.9998 | 10 |
| Rutin | 3.5-8000 | y=18915x + 298272 | 0.9998 | 3.5 |
| Peonidin 3-O-rutinoside | 5-900 | y=14348x + 35023 | 0.9997 | 5 |
| Strictosidine | 1.0-500 | y = 27175x + 11179 | 0.9998 | 1 |
| Catharanthine | 0.1-200 | y = 1362842x + 856727 | 0.9997 | 0.1 |
| Deacetylvindoline | 0.1-100 | y = 1076941x + 12409 | 1 | 0.1 |
| Vindoline | 0.1-300 | y = 1341203x + 707158 | 0.9995 | 0.1 |
| Vindorosine | 0.2-450 | y= 859082x + 1547496 | 0.9995 | 0.2 |
| Tabersonine | 0.1-250 | y = 928020x + 48163 | 0.9997 | 0.1 |
| Serpentine | 0.12-600 | y = 1382354x + 1665432 | 0.9993 | 0.12 |
| Vindolinine | 0.1-245 | y=1438007x + 1085239 | 0.9997 | 0.1 |
| Coronaridine | 0.1-200 | y = 1918865x + 663204 | 0.9999 | 0.1 |
| Anhydrovinblastine | 0.15-150 | y = 1509391x + 273636 | 0.9996 | 0.15 |
| Vinblastine | 0.06-100 | y = 1207682x + 444112 | 0.9991 | 0.06 |

<sup>a</sup>Each point of calibration curve was measured in triplicate.
<sup>b</sup>The LOQ was estimated in the lowest analyte concentration injected that yielded a signal-to-noise (S/N) ratio of $\geq 10$ in three replicates.

**Supplementary Table 5.** Summary of parameters of scMS for 5 tissues.

| Tissue of origin | No. of cells analyzed | No. of features with formula | No. of features with MS<sup>2</sup> data | No. of features/cell<sup>a,b</sup> |
| --- | --- | --- | --- | --- |
| Leaf | 202 | 557 | 295 | 17-119 |
| Root | 187 | 869 | 438 | 10-280 |
| Petal SA | 232 | 765 | 321 | 27-106 |
| Petal LBE | 241 | 1373 | 567 | 72-221 |
| Petal ABH | 244 | 513 | 239 | 28-140 |

<sup>a</sup>No. of features with formula assignment with error lower than 2 ppm and after blank filtering.
<sup>b</sup>10 cells/dataset were randomly selected for this calculation.

**Supplementary Table 6.** List of all chemicals used in this study.

| No. | Compound | CAS | Supplier | SKU/Code |
| --- | --- | --- | --- | --- |
| **Alkaloids** |  |  |  |  |
| 1 | Catharanthine | 2468-21-5 | Sigma Aldrich | SML0259 |
| 2 | Tabersonine | 4429-63-4 | Sigma Aldrich | SMB00452 |
| 3 | Vindoline | 2182-14-1 | Acros Organics BVBA | 462270010 |
| 4 | Anhydro Vinblastine disulfate | 81165-17-5 | TRC Canada | TRC-A659750 |
| 5 | Vinblastine sulfate | 143-67-9 | Acros Organics BVBA | 203450050 |
| 6 | Serpentine hydrogen tartrate | 58782-36-8 | Sequoia Research Products Ltd | SRP01300s |
| 7 | Strictosidine (10 mM) | 20824-29-7 | Synthesized |  |
| 8 | Ajmaline | 4360-12-7 | Extrasynthese | 0623 |
| 9 | Ajmalicine | 483-04-5 | Sigma Aldrich | 41111 |
| 10 | Coronaridine | 467-77-6 | Isolated |  |
| 11 | Deacetylvindoline | 3633-92-9 | TRC Canada | D288745 |
| 12 | Vindolinine | 5980-02-9 | Advanced ChemBlocks Inc | U105985 |
| 13 | Hörhammericine |  | Synthesized |  |
| 14 | Vindorosine | 5231-60-7 | Isolated |  |
| **Iridoids** |  |  |  |  |
| 1 | Secologanin | 19351-63-4 | Sigma Aldrich (SAFC) | 50741 |
| 2 | Loganic acid | 22255-40-9 | Extrasynthese | 0230 S |
| **Anthocyanin** |  |  |  |  |
| 1 | Petunidin-3-*O*-glucoside chloride | 6988-81-4 | Extrasynthese | 0919 |
| 2 | Peonidin-3-*O*-glucoside chloride | 6906-39-4 | Extrasynthese | 0929 S |
| 3 | Peonidin-3-*O*-rutinoside chloride | 27539-32-8 | Extrasynthese | 0945 |
| 4 | Hirsutidin chloride | 4092-66-4 | TRC Canada | TRC-H356800 |
| 5 | Peonidin chloride | 134-01-0 | Extrasynthese | 0906 S |
| **Flavonoids** |  |  |  |  |
| 1 | Rutin trihydrate | 250249-75-3 | Sigma Aldrich | 78095 |
| 2 | Mauritianin | 109008-28-8 | BIOMOL GmbH | 34326 |
| **Protoplast isolation** |  |  |  |  |
| 1 | Macerozyme R-10 from *Rhizopus* sp. Lyophil. | 9032-75-1 | SERVA | 28302.02 |
| 2 | Cellulase »Onozuka« R-10 aus<br>Trichoderma viride ca. 1 U/mg | 9012-54-8 | SERVA | 16419.03 |
| 3 | Pectinase from Aspergillus niger | 9032-75-1 | Sigma Aldrich | P4716 |
| 4 | Fluorescein Diacetate | 596-09-8 | TCI | F0240 |
| 5 | D-Mannitol | 69-65-8 | Sigma Aldrich | 6356 |
| 6 | MES hydrate | 1266615-59-1 | Sigma Aldrich | M8250 |
| 7 | Bovine serum albumin | 9048-46-8 | Sigma Aldrich | A2153 |
| **Sample preparation and mobile phases** |  |  |  |  |
| 1 | Formic Acid Optima LC/MS | 64-18-6 | Fisher Scientific | A117 |
| 2 | Acetonitrile OPTIMA® LC/MS<br>GRADE | 75-05-8 | Fisher Scientific | A955-212 |
| 3 | Methanol UHPLC-MS | 67-56-1 | Fisher Scientific | A458-1 |

**Supplementary Table 7.**
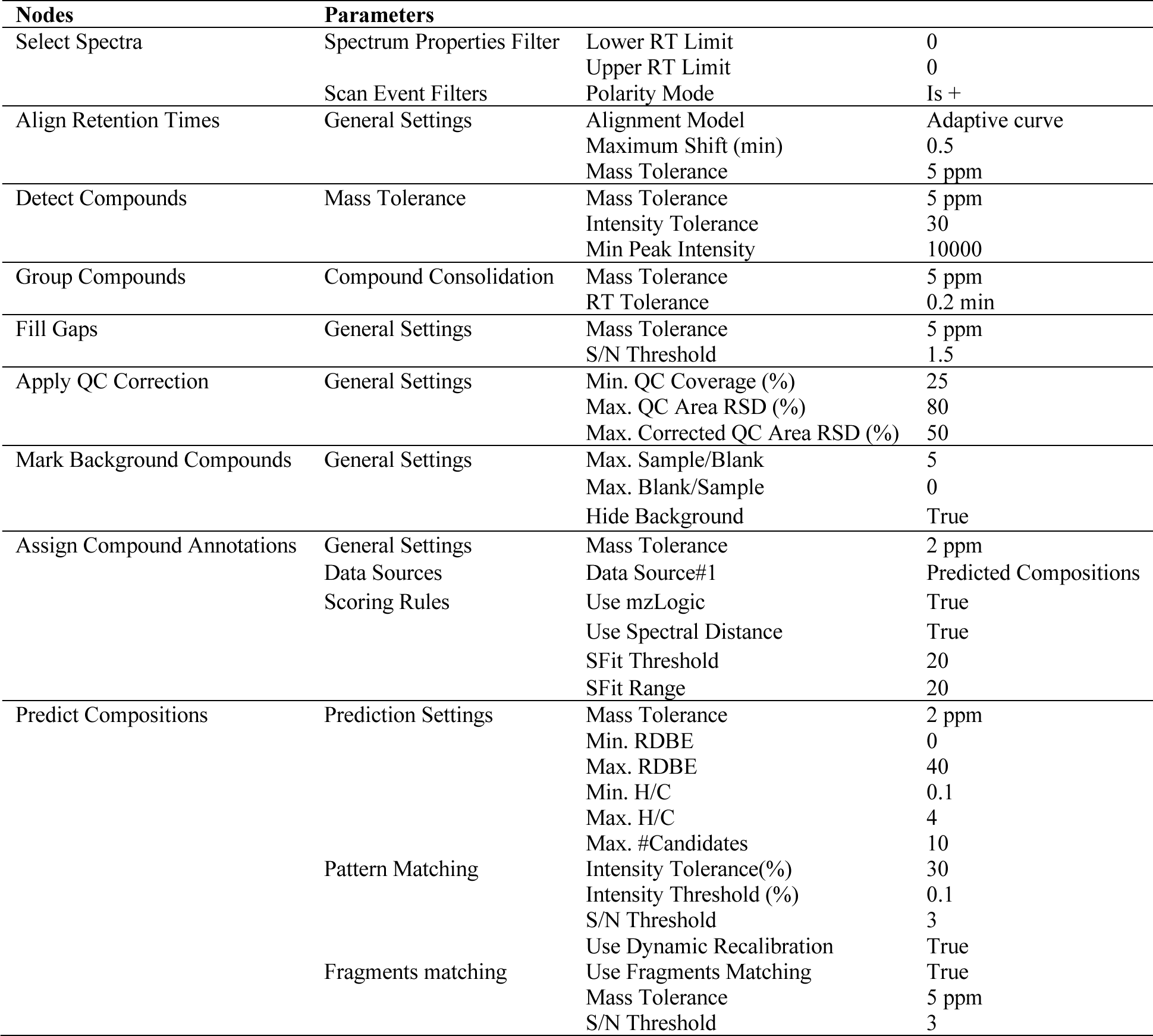
Compound Discoverer™ important parameters.

### Supplementary Data and Source File

**Supplementary Data 1.** Bulk untargeted analysis dataset with molecular formula assigned less than 2 ppm error (.xlsx file).

**Supplementary Data 2.** Feature classification to 3 main chemical classes using SIRIUS (.xlsx file).

**Supplementary Data 3.** Single cell untargeted analysis dataset with molecular formula assigned less than 2 ppm error (.xlsx files).

**Supplementary Data 4.** Quantification analysis dataset for single cells (.xlsx files).

**Supplementary Data 5.** Source data for Supplementary Fig. 18 (.xlsx file).

**Supplementary Data 6.** Source data for Supplementary Fig. 19 (.xlsx file).

**Source File 1.** Fragmentation data from all compounds identified in this study.

## References

1. Ozber, N. & Facchini, P.J. Phloem-specific localization of benzylisoquinoline alkaloid metabolism in opium poppy. Journal of Plant Physiology 271, 153641 (2022).

2. Lv, Q., Li, X., Fan, B., Zhu, C. & Chen, Z. The Cellular and Subcellular Organization of the Glucosinolate–Myrosinase System against Herbivores and Pathogens. International Journal of Molecular Sciences 23, 1577 (2022).

3. Kang, M., Choi, Y., Kim, H. & Kim, S.-G. Single-cell RNA-sequencing of Nicotiana attenuata corolla cells reveals the biosynthetic pathway of a floral scent. New Phytologist 234, 527–544 (2022).

4. Courdavault, V. et al. A look inside an alkaloid multisite plant: the Catharanthus logistics. Current Opinion in Plant Biology 19, 43–50 (2014).

5. Li, C. et al. Single-cell multi-omics in the medicinal plant Catharanthus roseus. Nature Chemical Biology 19, 1031–1041 (2023).

6. Sun, S. et al. Single-cell RNA sequencing provides a high-resolution roadmap for understanding the multicellular compartmentation of specialized metabolism. Nature Plants 9, 179–190 (2023).

7. de Souza, L.P., Borghi, M. & Fernie, A. Plant Single-Cell Metabolomics—Challenges and Perspectives. International Journal of Molecular Sciences 21, 8987 (2020).

8. Misra, B.B., Assmann, S.M. & Chen, S. Plant single-cell and single-cell-type metabolomics. Trends in Plant Science 19, 637–646 (2014).

9. Fujii, T. et al. Direct metabolomics for plant cells by live single-cell mass spectrometry. Nature Protocols 10, 1445–1456 (2015).

10. Yamamoto, K. et al. Cell-specific localization of alkaloids in Catharanthus roseus stem tissue measured with Imaging MS and Single-cell MS. Proc Natl Acad Sci U S A 113, 3891–3896 (2016).

11. Dührkop, K. et al. SIRIUS 4: a rapid tool for turning tandem mass spectra into metabolite structure information. Nat Methods 16, 299–302 (2019).

12. Dührkop, K. et al. Systematic classification of unknown metabolites using high-resolution fragmentation mass spectra. Nat Biotechnol 39, 462–471 (2021).

13. Kim, H.W. et al. NPClassifier: A Deep Neural Network-Based Structural Classification Tool for Natural Products. J Nat Prod 84, 2795–2807 (2021).

14. Forsyth, W.G.C. & Simmonds, N.W. Anthocyanidins of Lochnera rosea. Nature 180, 247–247 (1957).

15. Filippini, R., Caniato, R., Piovan, A. & Cappelletti, E.M. Production of anthocyanins by Catharanthus roseus. Fitoterapia 74, 62–67 (2003).

16. Xiao, Y., Tang, Y., Huang, X., Zeng, L. & Liao, Z. Integrated Transcriptomics and Metabolomics Analysis Reveal Anthocyanin Biosynthesis for Petal Color Formation in Catharanthus roseus. Agronomy 13, 2290 (2023).

17. Guedes, J.G. et al. The leaf idioblastome of the medicinal plant Catharanthus roseus is associated with stress resistance and alkaloid metabolism. J Exp Bot 75, 274–299 (2023).

18. Dai, Y. et al. in Advances in Botanical Research, Vol. 97. (eds. R. Verpoorte, G.-J. Witkamp & Y.H. Choi) 159–184 (Academic Press, 2021).

19. Choi, Y.H. et al. Are Natural Deep Eutectic Solvents the Missing Link in Understanding Cellular Metabolism and Physiology? Plant Physiol 156, 1701–1705 (2011).

20. Buhrman, K., Aravena-Calvo, J., Ross Zaulich, C., Hinz, K. & Laursen, T. Anthocyanic Vacuolar Inclusions: From Biosynthesis to Storage and Possible Applications. Front Chem 10 (2022).

21. Miettinen, K. et al. The seco-iridoid pathway from Catharanthus roseus. Nature Communications 5, 3606 (2014).

22. Mahroug, S., Courdavault, V., Thiersault, M., St-Pierre, B. & Burlat, V. Epidermis is a pivotal site of at least four secondary metabolic pathways in Catharanthus roseus aerial organs. Planta 223, 1191–1200 (2006).

23. Konno, K., Hirayama, C., Yasui, H. & Nakamura, M. Enzymatic activation of oleuropein: A protein crosslinker used as a chemical defense in the privet tree. Proceedings of the National Academy of Sciences 96, 9159–9164 (1999).

24. Larsen, B. et al. Identification of Iridoid Glucoside Transporters in Catharanthus roseus. Plant Cell Physiol 58, 1507–1518 (2017).

25. Kulagina, N., Méteignier, L.-V., Papon, N., O’Connor, S.E. & Courdavault, V. More than a Catharanthus plant: A multicellular and pluri-organelle alkaloid-producing factory. Current Opinion in Plant Biology 67, 102200 (2022).

26. O’Connor, S.E. & Maresh, J.J. Chemistry and biology of monoterpene indole alkaloid biosynthesis. Nat Prod Rep 23, 532–547 (2006).

27. Asma, S.T. et al. Natural Products/Bioactive Compounds as a Source of Anticancer Drugs. Cancers 14, 6203 (2022).

28. Zhang, J. et al. A microbial supply chain for production of the anti-cancer drug vinblastine. Nature 609, 341–347 (2022).

29. Koroleva, O.A., Gibson, T.M., Cramer, R. & Stain, C. Glucosinolate-accumulating S-cells in Arabidopsis leaves and flower stalks undergo programmed cell death at early stages of differentiation. The Plant Journal 64, 456–469 (2010).

30. Koroleva, O.A. et al. Identification of a New Glucosinolate-Rich Cell Type in Arabidopsis Flower Stalk. Plant Physiol 124, 599–608 (2000).

31. Contin, A., van der Heijden, R. & Verpoorte, R. Accumulation of loganin and secologanin in vacuoles from suspension cultured Catharanthus roseus cells. Plant Sci 147, 177–183 (1999).

32. Carqueijeiro, I., Noronha, H., Duarte, P., Gerós, H. & Sottomayor, M. Vacuolar Transport of the Medicinal Alkaloids from Catharanthus roseus Is Mediated by a Proton-Driven Antiport Plant Physiol 162, 1486–1496 (2013).

33. Yuan, C. & Yang, H. Research on K-Value Selection Method of K-Means Clustering Algorithm. J 2, 226–235 (2019).

34. Gu, Z. & Hübschmann, D. Make Interactive Complex Heatmaps in R. Bioinformatics 38, 1460–1462 (2021).

35. Clark, I.C. et al. Microfluidics-free single-cell genomics with templated emulsification. Nat Biotechnol 41, 1557–1566 (2023).

36. Kaminow, B., Yunusov, D. & Dobin, A. STARsolo: accurate, fast and versatile mapping/quantification of single-cell and single-nucleus RNA-seq data. bioRxiv, 2021.2005.2005.442755 (2021).

37. Yang, S. et al. Decontamination of ambient RNA in single-cell RNA-seq with DecontX. Genome Biology 21, 57 (2020).

38. Stuart, T. et al. Comprehensive Integration of Single-Cell Data. Cell 177, 1888–1902.e1821 (2019).

39. Kim, J.-Y. et al. Distinct identities of leaf phloem cells revealed by single cell transcriptomics. The Plant Cell 33, 511–530 (2021).

## References

1. Li, C. et al. Single-cell multi-omics in the medicinal plant Catharanthus roseus. Nature Chemical Biology 19, 1031–1041 (2023).

2. Yuan, C. & Yang, H. Research on K-Value Selection Method of K-Means Clustering Algorithm. J 2, 226–235 (2019).

